# Validation of diffusion and exchange imaging biomarkers via simultaneous real-time NMR and optical microscopy

**DOI:** 10.64898/2025.12.30.697003

**Authors:** Rea Ravin, Nathan H. Williamson, Teddy X. Cai, Natalia Gudino, Peter J. Basser

## Abstract

Diffusion MRI can reflect features of tissue microstructure and homeostasis, but direct validation in living neural tissue remains challenging. Here we combine optical microscopy and NMR for real-time recording on viable *ex vivo* neural tissue during perturbations. Simultaneous high-temporal-resolution NMR and optical microscopy are used to monitor apparent diffusion coefficient (ADC), apparent exchange rate (AXR), intrinsic optical signal (IOS), and intracellular calcium in *ex vivo* neonatal mouse spinal cord during osmotic and ionic perturbations. We find that ADC correlates strongly with IOS, while AXR decreases with metrics related to depolarization. ADC and AXR are sensitive to distinct features of cellular swelling, supporting their complementary roles in probing tissue viability and function.

---

Diffusion MRI can probe features of cellular structure and homeostasis in living tissues such as the central nervous system (CNS), but its various imaging biomarkers often lack direct cross-validation. ^1^ Most validation efforts have combined MRI with microscopy, but the approach and findings are not easily generalized due to their use of fixed *ex vivo* tissue, ^2–6^ where cellular function is halted and membrane permeability and compartmental volumes are altered. ^7,8^ While *in vivo* diffusion MRI has the potential to elucidate some physiological processes, ^9–14^ it suffers from low sensitivity, slow acquisition, confounding effects from blood flow and respiration, ^15,16^ and challenges when combined with other modalities. ^17–19^ We show that real-time NMR measurements with viable *ex vivo* tissue can overcome these limitations by preserving physiological function while enabling exquisite experimental control and relative ease to combine virtually any type of optical microscopy.

NMR can characterize diffusion and the steady-state exchange of water noninvasively. ^20,21^ Magnetic field gradients can be used to encode molecular displacements along the direction of the gradient and to measure the apparent diffusion coefficient (ADC). Differences in diffusion properties within the intracellular and extracellular environments can be observed on timescales where water, on average, has felt the effect of plasma membranes but has not yet permeated them. On these timescales, water inside cells is less mobile because it is bound and restricted by membranes, whereas water outside is more mobile because it can diffuse through a tortuous but connected extracellular space (ECS). In this way the ADC may be used as a proxy for cell volume, but not necessarily quantitatively due to structural heterogeneity. In particular, the heterogeneity of cell sizes and orientations of cellular processes can lead to water in some intracellular domains appearing more mobile than others. Exchange between environments where water mobility differs can be measured by encoding the same water at two instances separated by a mixing time where the water can communicate between environments. ^22,23^ In this way, the apparent exchange rate constant (AXR) is sensitive to membrane permeability, ^24,25^ however not quantitatively due to the heterogeneity in cell size, and thus cell surface-to-volume ratios, and not specifically due to the sensitivity to other exchange pathways. In particular for gray matter, diffusion-mediated, geometric exchange along branching cell processes, ^26,27^ between processes and soma, ^27,28^ between dendritic spines and shafts, ^29,30^ and localization-driven exchange between regions near and far from membranes ^31^ can occur in combination with transmembrane exchange, leading to multisite exchange. ^32,33^

Steady-state water exchange is a fundamental aspect of cellular homeostasis. ^34^ Water exchange occurs by passive diffusion ^32,35^ and, potentially, active transport processes. ^36–38^ NMR provides one of the only direct means to measure steady-state water exchange, but measurements are time consuming because they require repeated scans with different encodings. Validation of exchange metrics is especially challenging since exchange and structure are intertwined and there is no method for direct comparison. This necessitates approaches that characterize as many inputs as possible simultaneously and interpret them within the physiological context of the living system. In CNS gray matter, exchange across densely packed neuronal and glial processes is expected to be rapid due to their high surface-to-volume ratios ^39,40^ and potentially high membrane permeability. ^41,42^ Capturing these rapid dynamics demands methods with both the spatial resolution to probe fine structural compartments and the temporal resolution to follow fast exchange processes—capabilities achieved by the real-time NMR methods developed here.

In this study, we combine new real-time, high-temporal-resolution NMR with high-temporal resolution optical microscopy of intrinsic optical signals (IOS), ^43–51^ and intra-cellular calcium ([Ca^2+^]_i_) imaging to validate the sensitivity of the apparent diffusion coefficient (ADC) and apparent exchange rate (AXR) to changes in cellular structure and function. Using viable *ex vivo* neonatal mouse spinal cords with a solenoid RF coil provides a high filling factor and roughly a 10-fold SNR improvement over previous flat coil designs. ^52^ To accommodate an inverted microscope positioned beside the NMR magnet, an objective inverter was used. The single-sided permanent magnet provides access to the sample from above and provides an extremely strong *g* = 15.3 T/m static gradient (SG) for measuring ADCs on sub-millisecond diffusion timescales, setting the higher limit for measurable AXRs at 1000 s^−1^, an order of magnitude higher than can be measured with standard pulsed field gradient (PFG) approaches. During osmotic pressure (sucrose) and ionic (KCl) perturbations, we find that ADC correlates closely with IOS, suggesting similar biophysical origins—namely, cell shrinking and swelling. AXR remains relatively stable during periods of increased osmolarity but drops when [Ca^2+^]_i_ peaks during KCl addition, suggesting that it reflects loss of tissue homeostasis during spreading depolarization.

## Results

### Real-time diffusion and exchange measurement and validation

We first develop quantitative real-time NMR methods to characterize the homeostatic state fast enough to observe changes induced by osmotic (sucrose) and ionic (KCl and NaCl) perturbations to the media bathing the spinal cord. Concentration gradients across the ≈ 0.7 mm radius of tissue are expected to equilibrate on a timescale of ≈ 1 minute for water, ^8^ slightly longer for K^+^ and Na^+^, and roughly four times longer for sucrose based on their self-diffusion coefficients. ^53–55^ Mechanisms such as volume regulation and K^+^ buffering act to preserve homeostasis during perturbations, but at times these are not enough. Excess extracellular K^+^ triggers spreading depolarization (SD)—an “all-or-none” cascade of neuronal firing and K^+^ release propagating at millimeters per minute. ^56^ SD occurs across a range of pathologies, including migraine aura, traumatic brain injury, and stroke. ^57^ Bath application of 50 mM KCl is expected to evoke SD at the cord surface, spreading to the center within a minute without recovery, causing [Ca^2+^]_i_ elevation, and IOS, ADC and AXR reduction from cellular swelling and extracellular shrinkage. ^10,32,56,58,59^ To better match those dynamics—and to narrow the gap with the temporal resolution of microscopy—we developed a protocol which takes 40 seconds total per ADC and AXR measurement. This was achieved by isolating the experimental weighting parameter needed for the measurement.

The ADC measurement is based on SG spin echo (SG SE) diffusion encoding where the diffusion weighting (DW) is summarized by *b* = 2*/*3*γ*^2^*g*^2^*τ* ^3^ with gyromagnetic ratio *γ* and diffusion encoding time *τ* in the equation *S*(*b*) = *S*_0_ exp(−*b*ADC). With two free parameters, the analytical expression for finding the (1-dimensional) ADC with two signals *S*(1) and *S*(2) acquired at *b*(1) near zero and *b*(2) near 1*/*ADC is

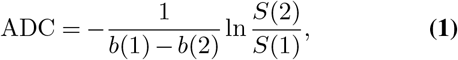

as is well-known in the field of diffusion MRI. ^60^

The AXR measurement is based on the SG diffusion exchange spectroscopy (DEXSY) experiment which applies two diffusion encodings (*b*_1_ and *b*_2_) separated by a mixing time (*t*_*m*_). ^8,22^ Early DEXSY implementations required hours to days per dataset because of extensive phase cycling and dense sampling of *b*_1_ and *b*_2_ values. ^8,22,61,62^ Later methods such as filter exchange spectroscopy and imaging (FEXSY, FEXI) reduced acquisition time by optimizing signal sampling. ^23,63–65^ Further improvements came from understanding how to isolate diffusion, exchange, and relaxation weightings. ^66–72^

For two-site, barrier-limited exchange—where water molecules traverse the intracellular space (environment *i*) many times before crossing to the extracellular space (*o*)—exchange-weighted DEXSY signals attenuate with the apparent exchange rate (AXR) and longitudinal relaxation rate (*R*_1_) according to

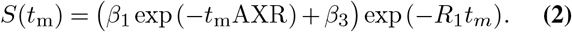

In this case, AXR = *k* = *k*_*io*_ + *k*_*oi*_, the sum of the rate constants for exchange between the two environments. The diffusion exchange ratio (DEXR) method uses prior knowledge of *R*_1_ to reduce Eq. 2 to a three-parameter model. ^68,72^ This allows AXR estimation from three relaxation-adjusted signals—acquired at short (*t*_*m*,short_ ≪ 1*/*AXR), intermediate (*t*_*m*,int_ ≈ ≫ 1*/*AXR), and long (*t*_*m*,long_ ≫ 1*/*AXR) mixing times via

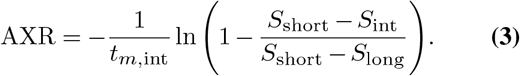

Both SE and DEXSY MR sequences were repeated four times per signal to cover the phase cycle steps which add together desired signals and subtract unwanted ones, ^73,74^ still with a substantial improvement in efficiency compared to earlier static-gradient DEXSY implementations. ^8,62^ With a 2 second repetition time, this leads to 16 seconds for an ADC measurement and 24 second for an AXR measurement, which to our knowledge is the fastest NMR-based exchange measurement to-date. These sequences acquire signals using a Carr–Purcell–Meiboom–Gill (CPMG) echo train, ^75–77^ which simultaneously yields DW and transverse relaxation rates (*R*_2_). While this ultra-fast approach cannot identify and characterize multiple exchanging and non-exchanging compartments, it also obviates the assumptions and complexity of models in which at least one of those compartments is restricted.

The 3-point AXR measurement has never been used before. We first validated it using both numerical simulations and subsampled DEXSY data. ^34^ Simulations of two-site exchange showed quantitative accuracy for true rate constants (*k*) between roughly 20 and 150 s^−1^ (Fig. S1), consistent with the range we expect from previous studies with longer recordings. ^34^ The range of accuracy could be tuned by shifting the mixing time values (Fig. S2). In data subsampled from oxygen and glucose deprivation (OGD) experiments on *ex vivo* neonatal mouse spinal cords, the method remained quantitative under normal conditions (AXR ≈ 140 s^−1^) but overestimated exchange after OGD, when AXR dropped to ≈ 40 s^−1^ (Fig. S3). This overestimation likely arises because AXR is sensitive to mixing-time placement in systems exhibiting multiexponential geometric exchange. ^27,74,78,79^

### Experimental setup

Simultaneous NMR and microscopy was achieved using a low-field single-sided magnet projecting a magnetic field from below while the microscope viewed the sample from above (Fig. 1). A custom test chamber and RF probe with a solenoid coil were used to maintain tissue viability while maximizing filling factor and SNR. ^8,34^ The sample was viewed through a 650 *µ*m gap in the solenoid coil. An objective inverter permitted the use of an inverted microscope.

**Fig. 1.**
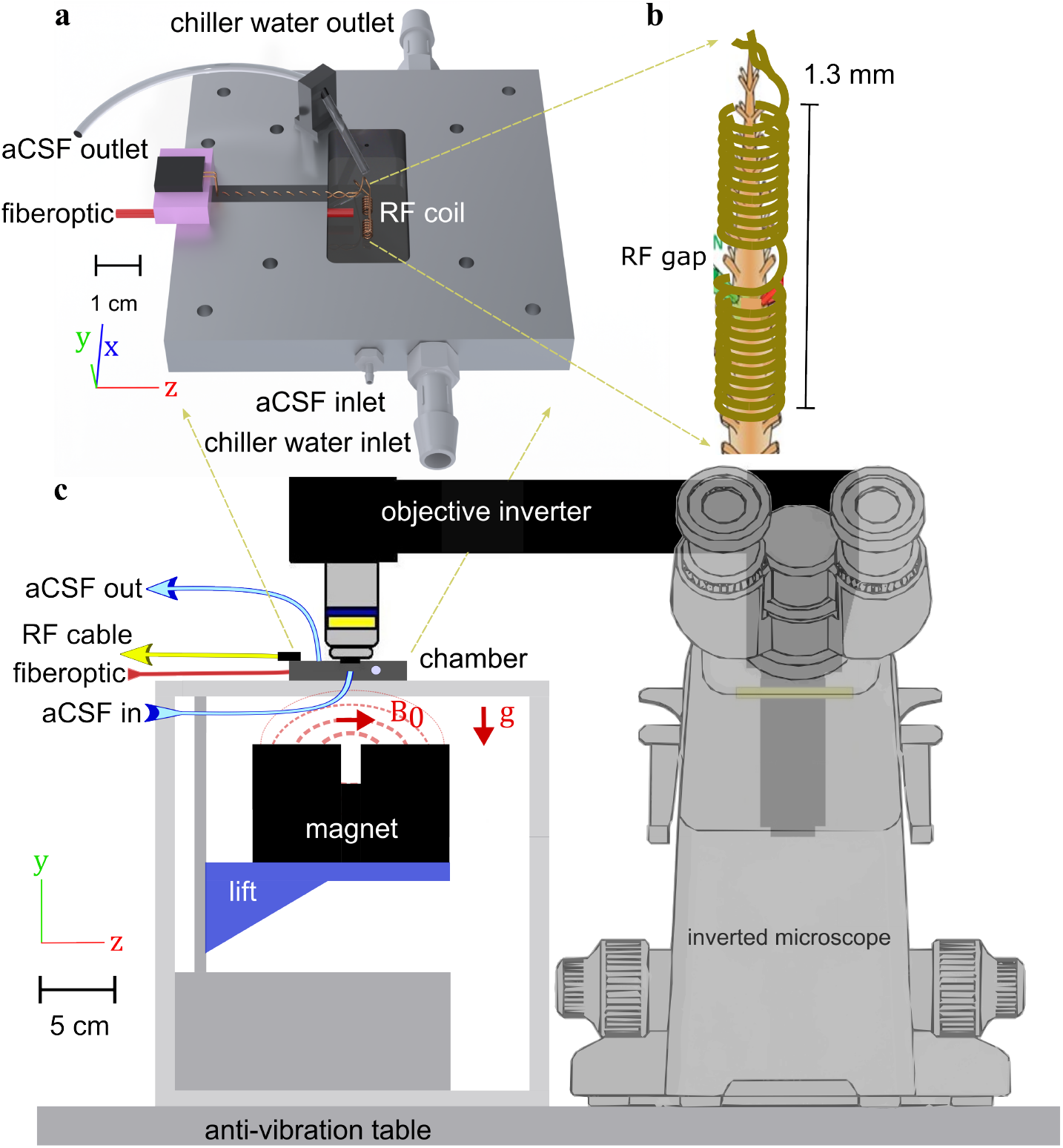
Experimental setup. a) 3-D technical drawing of the test chamber. b) Drawing of the solenoid RF coil containing a mouse spinal cord showing the gap in the RF coil where the sample is imaged. c) Technical drawing of the experimental setup showing the single sided permanent magnet projecting a magnetic field on the sample from below and the inverted microscope imaging the sample from above.

### Intrinsic Optical Signal: Comparison of reflected and transmitted light

On the microscopy side, we focused on intrinsic optical signal (IOS) imaging because, while the mechanism is completely different and relies on tissue light scattering or transmittance changes, like diffusion MRI, it is sensitive to cellular-scale microstructural changes averaged over many cells in *ex vivo* CNS tissue. ^43–51^ Although IOS is relatively simple to acquire, it has not previously been performed simultaneously with NMR.

IOS contrast depends on the angle between the light source and the objective. ^45^ The single-sided magnet is sitting below the stage in close proximity to the stage and chamber, making it difficult to orient the light source below the sample. For this reason, we decided to use a 90^◦^ configuration, in which the fiberoptic light source and objective are oriented perpendicularly (Fig. 1). Then, the detected light primarily arises from reflection and scattering within the sample. In contrast, cylindrical, closed-bore magnets could more easily support a 180^◦^ configuration, where transmitted rather than reflected light reaches the detector. Because transmittance is expected to vary inversely with reflectance during microstructural changes such as cell swelling or shrinkage, ^45^ we compared both geometries (IOS_90_ and IOS_180_) in separate experiments using the same system, but the latter performed without NMR.

Following the addition of 50 mM KCl for 20 minutes and subsequent washout, IOS_90_ decreased while IOS_180_ increased (Fig. 2a) as expected from previous studies. ^45^ When normalized to their peaks the signals were highly similar (Fig. 2b), consistent with prior findings that both reflected and transmitted light track cellular swelling. ^45^ For compatibility with the NMR setup, subsequent experiments used only IOS_90_ (hereafter referred to simply as IOS).

**Fig. 2.**
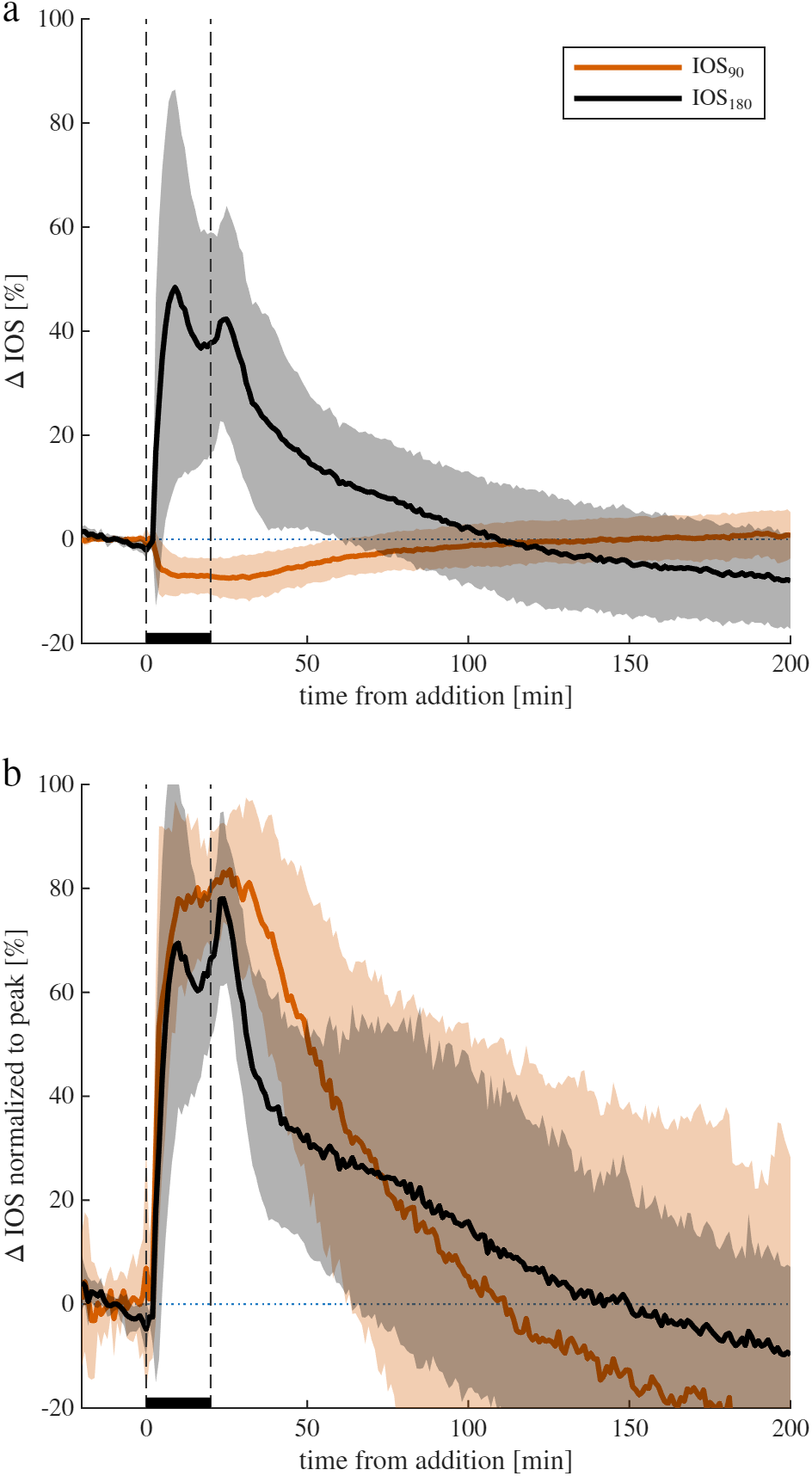
Comparison of 90^◦^ and 180^◦^ configurations. a) Percent change in IOS from baseline (mean ± standard deviation) for reflected light (IOS_90_, n = 13) and transmitted light (IOS_180_, n = 9) setups during exposure to 50 mM KCl at *t* = 0 and washout after 20 min. b) Normalized signals scaled to the minimum (IOS_90_) or maximum (IOS_180_) values.

### Simultaneous real-time NMR and microscopy demonstration

Simultaneous microscopy and NMR were then initiated with real-time recordings. We used microscopy to do simultaneous imaging of [Ca^2+^]_i_ and IOS. Spinal cords were bolus-injected with fluorescent indicator Rhod-3 AM for [Ca^2+^]_i_ imaging, and baseline NMR recordings were performed with the standard longer DEXR, *T*_1_, and diffusion measurements. This served as a quality control on the viability of the sample, as AXR provides an absolute value that can be compared between samples. ^34^ Samples for which AXR started too low (below 100 s^−1^) were discarded because they were found previously to run down and not recover. ^34^

Sample recordings from a 50 mM KCl perturbation experiment are shown in Fig. 3. Raw NMR signals from two SG SE and three SG DEXSY acquisitions were processed into ADC and AXR time-series using Eqs. 1 and 3, respectively (Fig. 3a–d). Previous studies have shown that baseline ADC values vary across samples ^34^. Because of this, subsequent figures report percent changes rather than absolute values. In contrast, baseline AXR values show little variation between samples, consistent with the interpretation that absolute AXR reflects tissue viability. ^34^

**Fig. 3.**
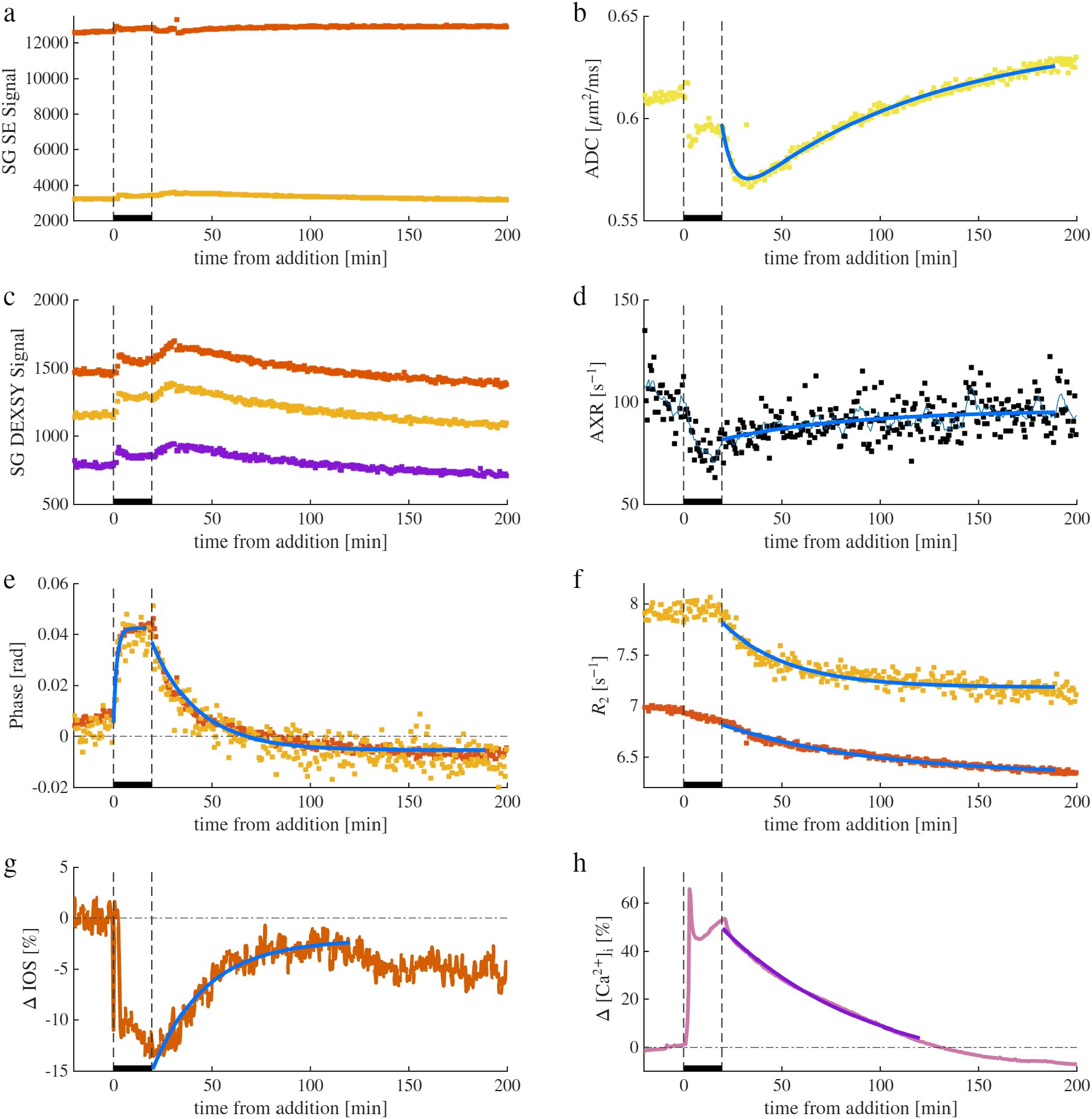
Example real-time NMR and microscopy recording during an experiment involving 50 mM KCl addition and washout. real-time recordings of two (*b* = 0.025 and 2.25 ms*/µ*m^2^, red and orange dots, respectively) raw spin echo diffusion signals (a), processed into one ADC measurement using Eq. 1 (b), and three (*t*_*m*_ = 0.02, 10, and 160 ms, red, orange, and blue dots, respectively) raw exchange-weighted DEXSY signals (c), processed into one AXR measurement using Eq. 3 (d). In (d), a 6-point moving average of AXR is also shown (solid blue line). e,f) Mean echo phase (e) and *R*_2_ (f) recorded using the spin echo sequence with *b* = 0.025 and 2.25 ms*/µ*m^2^ (red and orange dots, respectively). IOS images (g) and fluorescence images of [Ca^2+^]_i_ from the fluorescent indicator, Rhod-3 AM (h) are acquired simultaneously with the NMR, and the percent change in image intensity from a region of interest (ROI) is displayed. solid blue lines are mono- or bi- (for ADC) exponential fits, shown to guide the eye.

Each NMR signal represents the summed echoes of the CPMG acquisition block. From these, we also extract phase and transverse relaxation rates (*R*_2_) (Fig. 3e–i). Supplementary experiments on pure aCSF perturbed with 50 mM KCl show that phase is linearly related to the coil tuning frequency (Fig. S4). Tuning frequency shifts with sample impedance and in this way reflects sample dielectric properties (i.e., permittivity and electrical conductivity) ^80,81^. During the addition and washout of KCl, phase time-courses are used to measure the timescale of equilibration between the bathing medium and tissue (Fig. S5). Phase equilibrates with an exponential time constant of 1.2*±*0.2 min after 50mM KCl addition, and 9.0*±*2.7 min after washing to new media. Time constants are not significantly different when no sample is present, when NaCl is added instead of KCl, and when KCl is added 1 hr after 100 *µ*M of the Na^+^*/*K^+^–ATPase, ouabain was added earlier in the experiment (one-way ANOVA). This indicates that ion concentrations equilibrate in the sample roughly on the timescale of chamber turnover, independent of sample physiology. The absence of *b*-dependence in phase confirms negligible coherent flow along the gradient direction, as expected. ^82^

*R*_2_ reflects rotational mobility, ^83^ and increases with decreasing mobility. ^84^ Here, nearly all the signal comes from protons on water, ^8^ and *R*_2_ varies due to chemical and physical interactions between molecules and surfaces that affect water’s molecular tumbling rate. *R*_2_ and ADC are expected to be negatively correlated for diffusion in porous media because both are sensitive to the size of restrictions. ^85^ We find that *R*_2_ values increase with diffusion weighting, consistent with faster relaxation in more restricted environments. Their time courses differ: *R*_2_ declines steadily after KCl addition and washout, whereas diffusion-weighted *R*_2_ decreases only during washout. NMR metrics are acquired simultaneously with IOS and [Ca^2+^]_i_ (Fig. 3 g and h) (Fig. 3g,h). During the wash, the fluorescent signal related to [Ca^2+^]_i_ drops below baseline, likely due to the calcium indicator slowly diffusing out of the region of interest and not indicating [Ca^2+^]_i_ being lower than at baseline. Further trends will be discussed below. An example recording from a 100 mM sucrose perturbation experiment is shown in Fig. S6.

### Simultaneous real-time NMR and microscopy shows opposite effects of osmolytes and salts

In the first set of experiments, ADC and AXR from NMR were acquired simultaneously with IOS from microscopy during perturbations with 100 mM sucrose over 40 minutes (Fig. 4). Sucrose is an osmolyte that does not penetrate across the plasma membrane. The 100 mOsm osmotic pressure pulls water from the intracellular space (ICS) to the extracellular space (ECS), shrinking the cells until the intracellular pressure matches the extracellular pressure. This effect is seen by the increase in ADC and IOS. Volume regulation by partitioning additional ions and metabolites into the ICS subdues the effect. ^86^ AXR increases significantly (*p <* 0.001), but the effect is small (8.2*±*23.2%). Results of previous studies involving Na^+^*/*K^+^–ATPase inhibition and energy failure showed AXR remaining stable and dropping when homeostasis was lost, suggesting that provides a proxy for homeostasis. ^34^ Digging deeper into this mechanism, we found that water does not actively cycle with ions in this system. We did find that AXR is sensitive to tonicity and proposed a multi-site exchange mechanism in which AXR increases with ECS fraction due to biasing of the measurement from slower geometric exchange to faster transmembrane exchange. ^32,33^ The small effect of 100 mOsm sucrose on AXR and ADC has been shown previously by us ^32^ and is consistent with the multisite exchange mechanism and a slight increase in ECS fraction from an already significant level having little additional effect since transmembrane exchange already dominates.

**Fig. 4.**
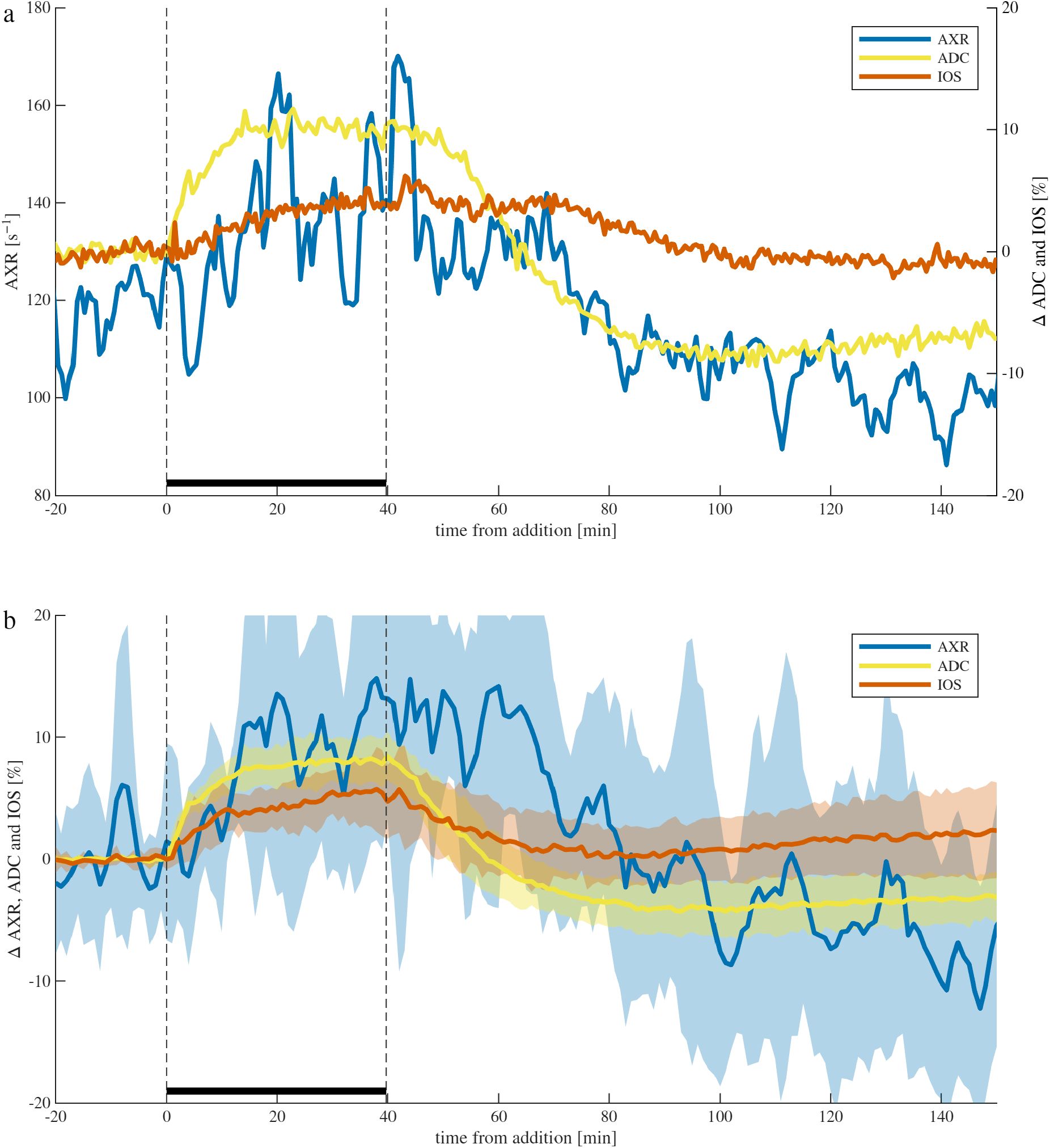
Simultaneous real-time NMR and microscopy shows cellular shrinkage caused by adding an osmolyte. a) A representative sample and b) means (lines) and standard deviations (shaded bands) across n=8 samples of real-time AXR (6-point moving average), ADC, and IOS recording during experiments involving 100 mM sucrose being added for 40 minutes and then washed away. *R*_2_ is presented in Fig. S7.

During the wash back to normal media, ADC and IOS recovered, although with slightly different timecourses (Fig. 4b). ADC decreased faster than IOS and overshot the baseline. The recovery from ADC overshoot was incomplete, with ADC values between 100 to 200 minutes after addition being 2.8*±*1.6% lower than baseline. The overshoot, indicative of cellular swelling, could be due to cells adjusting to a new steady state during the sucrose addition, but then having a higher intracellular osmolarity relative to the ECS, pulling water into the cells during the wash. AXR decreased significantly during the wash (*p <* 0.001), with AXR from 100 to 200 minutes after addition being 7*±*19% lower than baseline.

Out of the eight experiments, from the data acquired between *t* = 0 to 80 minutes after osmolyte addition, the correlation between AXR and ADC was only significant (*p <* 0.01) in two cases. The correlation between AXR and IOS was only significant in one case. The correlation between ADC and IOS was significant in six cases, and the correlation coefficient (*cc*) was 0.78*±*0.10 for those cases.

In the next set of experiments during 20 minute perturbations with 50 mM KCl, real-time NMR and microscopy data was acquired as previous but with [Ca^2+^]_i_ signal as well (Fig. 5). 50 mM KCl dissociates into 100 mM of ions, but its effect is unlike the 100 mM osmotic perturbation discussed above, first because cell membranes are permeable to ions. More importantly, 50 mM KCl initiates spreading depolarization, causing cellular swelling and ECS shrinkage. ^56^ Spreading depolarization is known to be accompanied by increased [Ca^2+^]_i_ which may be due to reversal of the Na^+^*/*Ca^2+^ exchanger, ^87^ activation of calcium channels, ^88^ mitochondrial depolarization and activation of the permeability transition pore, ^59^ transport from endoplasmic reticulum, ^89^ and transport through glial gap junctions. ^58^ While [Ca^2+^]_i_ imaging can be used to monitor SD, ^58^ the effect is indirect since SD can occur even when calcium is removed. ^56,90^ IOS is also used to monitor SD. ^58,59,90^ Interestingly, calcium waves have been found to occur before changes in IOS. ^90^ Since the KCl is bath-applied, the sample is expected to remain depolarized until the wash. Depolarization and loss of homeostasis is indicated by the [Ca^2+^]_i_ spike occurring 3 minutes into the perturbation (Fig S8). At the same time, swelling is indicated by the ADC and IOS decreasing to a minimum at 5 to 7 minutes, with ADC being 3.6*±*0.4 % reduced at 6 minutes. AXR also drops rapidly, by 21*±*3 % at 6 minutes, but then continues to drop at a slower rate, reaching a minimum of 26*±*5% at 14 minutes. While it appears that AXR was recovering at the end of the 20 minutes of high KCl, no recovery was seen in the presence of high KCl in experiments where the duration was increased to 40 minutes. In the multisite exchange mechanism, AXR can decrease with ECS volume fraction due to reduced sensitivity to faster transmembrane exchange and increased sensitivity to slower geometric exchange. ^33^ While the effect was smaller, interestingly *R*_2_ and DW *R*_2_ changed in the same direction as during the addition of 100 mM sucrose, with *R*_2_ decreasing to a minimum of −0.87*±*0.94 % at 2 minutes and DW *R*_2_ increasing to a maximum of 1.5*±*1.1 % at 16 minutes (Fig. S9). This indicates that *R*_2_ is not only affected by the structural length scale changes and is likely being affected by chemical changes as well here.

**Fig. 5.**
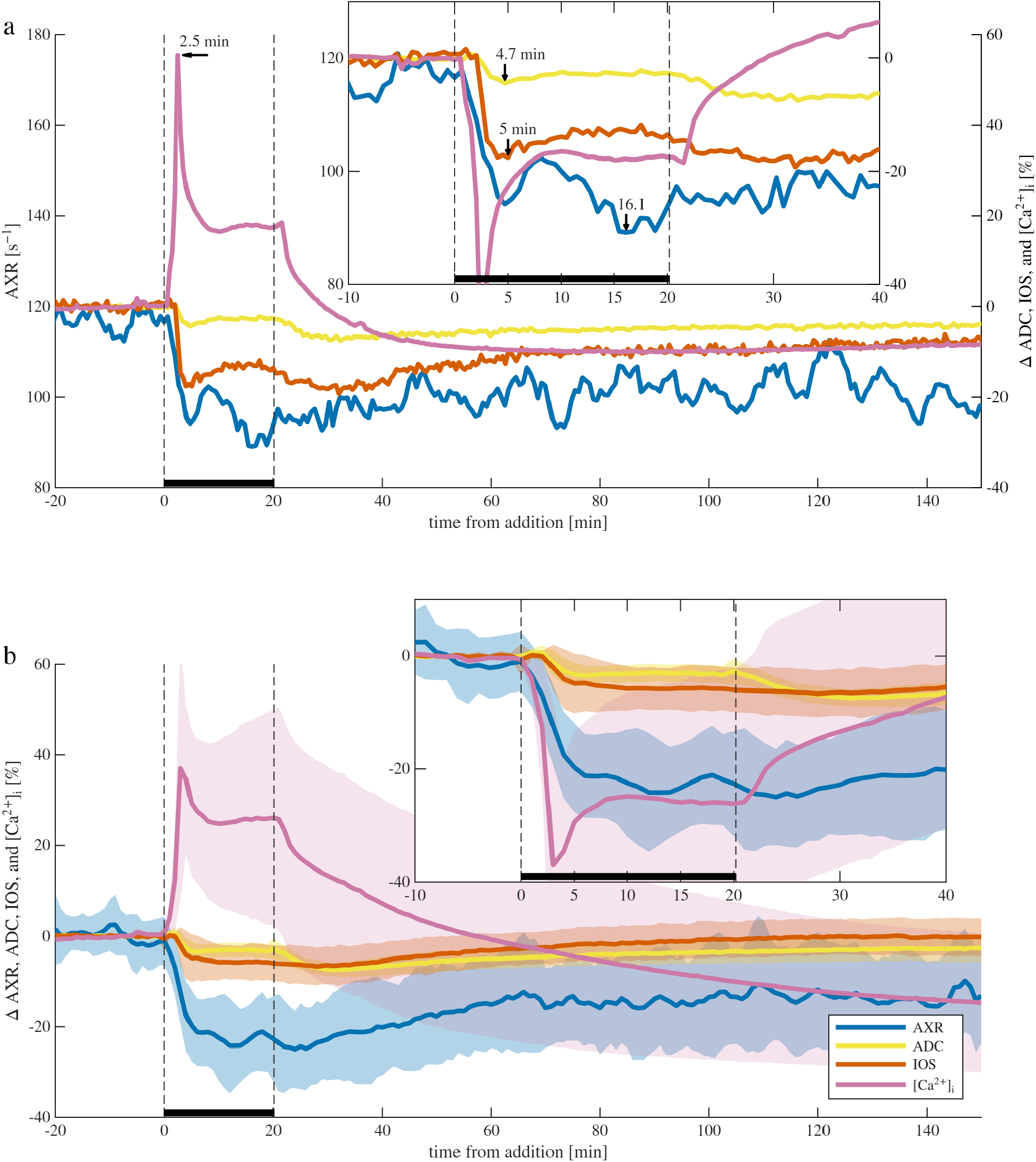
Simultaneous real-time NMR and microscopy reveal cellular swelling during depolarization from 50 mM KCl. a) Representative sample and b) mean ± SD of AXR (6-pt moving average), ADC, IOS, and [Ca^2+^]_i_ during 20 min KCl exposure and washout (*n* = 12, three of which included [Ca^2+^]_i_ imaging). Insets show zoomed regions with [Ca^2+^]_i_ inverted for comparison. Arrows in (a) mark signal minima and maxima. *R*_2_ is presented in Fig. S9.

When washing back to normal media, [Ca^2+^]_i_ quickly starts recovering, indicating calcium being transported out of the cells likely through normal operation of the Na^+^*/*Ca^2+^ exchanger. ^87^ Early in the wash the ADC and IOS decrease again prior to recovering. This may be partially because the built-up intracellular K^+^ accompanied by Cl^−^, particularly in astrocytes, ^91^ takes time to be pumped out but has an immediate osmotic swelling effect. The AXR, *R*_2_, and DW *R*_2_ do not completely recover. With these metrics being reduced, likely the recovery of ADC and IOS is not indicating a return to the same microstructural state.

Out of the 16 experiments, from the data acquired between *t* = 0 to 80 minutes after KCl addition, the correlation between AXR and ADC was significant in 12 cases (*cc* = 0.50*±*0.32). The correlation between AXR and IOS was significant in all case (*cc* = 0.57*±*0.15). The correlation between ADC and IOS was significant in 12 cases (*cc* = 0.58 *±* 0.21).

## Discussion

In this paper, we establish simultaneous real-time NMR and optical microscopy as a platform for translating knowledge from cellular neurophysiology to quantitative MRI and for validating the sensitivity of diffusion–exchange biomarkers to physiological and pathological processes. The temporal resolution of ADC and AXR measurements was pushed near its limit, with ADC and AXR requiring 16 seconds and 24 seconds to acquire, respectively. To the best of our knowledge, this is the highest temporal resolution yet reported for water exchange measurements. This made it possible to directly relate diffusion and exchange MRI biomarkers to rapid cellular physiological processes during osmotic and ionic perturbations.

The static gradient (SG) spin echo (SE) diffusion NMR experiments measured diffusivity on sub-millisecond timescales. With the large 15.3 T/m gradient of the NMR MOUSE, this measurement is highly sensitive to restriction on a length scale of 800 nm. ^92^ The DEXSY NMR experiment builds on this, measuring exchange between more and less restricted local environments during the prescribed mixing time. Previous studies have indicated that membranes are the sole source of restriction in the tissue. ^8^ Further, we found that DEXSY measurements are sensitive to transmembrane exchange, but they are also sensitive to other exchange pathways, in particular geometric exchange between more and less restricted environments in the cell. We showed that with passive exchange between multiple sites, the AXR could vary drastically depending on the ECS fraction. ^33^ We therefore have the working hypothesis that the AXR varies primarily with local ECS fraction. That said, we repeatedly have seen that it contains unique information distinct from the ADC. ^32,34^ This study continues to advance our understanding of the determinants of changes in the ADC and AXR in live tissue, with help from complementary optical measurements.

ADC and IOS were significantly correlated in 6/8 of osmotic perturbation experiments and 9/12 of KCl perturbation experiments. Similarities in the ADC and IOS timeseries data suggest similar biophysical origins, namely cellular swelling and shrinking. The differences observed, e.g., in the kinetics, could simply be due to NMR obtaining signal from a 400 nm slice through the length of the spinal cord, compared to the microscopy signal being obtained from a small field of view. While separate accounts in the optical microscopy ^43–51^ and diffusion MRI ^9,32,93–97^ literature independently suggest that both IOS and ADC are sensitive to cellular swelling, to the best of our knowledge this is the first time that they were simultaneously measured and their correlation has been reported. This suggests that physiological or pathological processes previously elucidated with IOS might also be visible in vivo with diffusion MRI.

During the osmotic perturbations, AXR was significantly correlated with ADC in only 2/8 experiments and with IOS in 1/8 experiments. During KCl perturbations, the correlations were stronger, with significant correlations between AXR and ADC in 12/16 experiments and between AXR and IOS in 16/16 experiments. This weak coupling during osmotic cell shrinkage and stronger coupling during pathological swelling is consistent with our previous observation of a sigmoidal relationship between AXR and ADC. ^32^

The multisite exchange mechanism predicts that both ADC and AXR depend on the extracellular space (ECS) volume fraction, but in fundamentally different ways. Whereas ADC reflects the volume-weighted average diffusivity across all compartments, AXR depends on the compartment volume fractions together with the eigenvalues of the exchange matrix. ^33^ Furthermore, compartments that exchange too slowly to contribute on the timescale of the measurement influence ADC but not AXR. ^98^ The remaining discrepancies between the three-site exchange model and experiment likely arise because neural tissue contains many more than three exchanging compartments whose volumes change heterogeneously during perturbation. Importantly, this more complex dependence means that AXR is sensitive to tissue homeostasis rather than cell volume alone, making it a promising marker of the homeostatic state of the tissue.

## Conclusion

While MRI and microscopy have each advanced our understanding of brain structure-–function relationships, studies that apply both methods simultaneously to viable tissue remain rare. The two techniques have largely complementary strengths with limited overlap, except in their shared sensitivity to cellular-scale tissue microstructure. Here, we demonstrate the direct benefits of combining them. For the first time, we show that ADC and IOS signals tend to track one another, indicating similar sensitivity to physiological and pathological cell swelling and shrinking. As a result, the IOS literature may help define mechanisms, limitations, and disease relevance of diffusion MRI. In contrast, AXR shows little correlation with ADC or IOS during osmolyte addition, when homeostasis is expected to be preserved, but exhibits stronger correlation during KCl addition, where we observe signs of depolarization and loss of homeostasis. This suggests that AXR is sensitive to the tissue’s homeostatic state and supports further investigation of its potential as a biomarker of tissue viability. Overall, these results illustrate how simultaneous microscopy and NMR can facilitate knowledge transfer between fields and accelerate progress at their intersection.

## Methods

### Ethics statement for animal experimentation

All experiments were carried out in compliance with the *Eunice Kennedy Shriver* National Institute of Child Health and Human Development Animal Care and Use Committee under Animal Study Proposal (ASP) # 21-025.

Details of NMR and tissue preparation methods can be found in Refs. 8,34,68, and 99, and are summarized briefly here.

### Mouse spinal cord sample preparation and experimental conditions

Spinal cords were isolated from postnatal day 1–4 Swiss Webster mice (Taconic Biosciences, Rensselaer, NY, USA) in low-calcium, high-magnesium artificial cerebrospinal fluid (aCSF) (128.35 mM NaCl, 4 mM KCl, 0.5 mM CaCl_2_, 6 mM MgSO_4_, 0.58 mM NaH_2_PO_4_, 21 mM NaHCO_3_, 30 mM D-glucose), bubbled with 95% O_2_/5% CO_2_). ^34^ After dissection, cords were transferred to the NMR chamber with the ventral side up, in standard aCSF (128.35 mM NaCl, 4 mM KCl, 1.5 mM CaCl_2_, 1 mM MgSO_4_, 0.58 mM NaH_2_PO_4_, 21 mM NaHCO_3_, 30 mM D-glucose) perfused at 7 mL/min and bubbled with 95% O_2_/5% CO_2_. Perturbations were introduced by adding sucrose or KCl from a stock solution directly to the bubbled reservoir, after which the tissue was washed in an open loop using 100 mL of fresh media. Temperature was maintained at 25*±*0.2^◦^C using a fiberoptic sensor (PicoM, Opsens Solutions Inc., Québec, Canada) and a chiller that circulated water through the aluminum block of the sample chamber (see Fig. 1) and through a heat exchanger jacketing the aCSF tubing upstream of the chamber.

### NMR experimental protocol and analysis methods

NMR experiments were performed with a single-sided permanent magnet (PM10 NMR MOUSE, Magritek, Aachen, Germany) at B_0_ = 0.3239 T and gradient strength *g* = 15.3 T*/*m (Fig. 1). ^100^ The static gradient was oriented in the *y*-direction, perpendicular to the spinal cord, defining the slice (400 *µ*m along the cord) and diffusion encoding direction.

Sample viability was first confirmed using diffusion, Diffusion Exchange Ratio (DEXR), and *T*_1_ recovery measurements following established protocols. ^34,72^

Subsequently, two-point ADC and three-point DEXR measurements were performed in parallel with microscopy image acquisition. The NMR RF pulse lengths were about 2 *µ*s. NMR pulse sequences consist of an encoding block and a CPMG acquisition block. ^75–77^ The acquisition block used 8000 echoes with 23.5, 25 or 28 *µ*s echo time, 4 *µ*s acquisition time, and 0.5 *µ*s dwell time and was analyzed in three ways. First, the mean angle between the real and imaginary components of the echoes yielded the phase. Second, the CPMG trains were fit with a single exponential for an *R*_2_ estimate weighted by the encoding block. Third, the real component of echoes 5 to 8000 was summed for an SE or DEXSY signal to be used in the measurement. The two-point ADC measurement used a static gradient spin echo (SE) encoding block ^73^ with *b* = 0.025 and 2.25 ms*/µ*m^2^ (*τ* = 0.131 and 0.587 ms) where *b* = 2*/*3*γ*^2^*g*^2^*τ*^3^.^101^ Points acquired immediately after RF tuning were omitted due to magnetization not being at steady state. ADC was then calculated from the two SE signals using Eq. 1. The three-point DEXR measurement used an exchange-weighted static gradient spin echo DEXSY encoding block ^68^ with (*b*_1_, *b*_2_) = (2.320, 2.170) ms*/µ*m^2^ (*τ*_1_, *τ*_2_ = 0.593, 0.581 ms) and mixing times *t*_*m*_ = 0.2, 10, and 160 ms, chosen to bracket expected apparent exchange rates (AXR ~50–150 s^−1^). DEXSY signals were corrected for relaxation by dividing by exp(−*t*_*m*_*R*_1_), where *R*_1_ = 1.5 s^−1^ was obtained from standard DEXR under baseline conditions. The relaxation-adjusted signals *S*_short_, *S*_int_, and *S*_long_ were then combined to estimate the apparent exchange rate constant via Eq. 3. With 2 s repetition time, 4 scans per signal (for phase cycles) and 5 signals total, the total acquisition time was 40 seconds per ADC and AXR measurement.

### Microscopy

An Axiovert 200 M wide-field inverted microscope (Zeiss, Germany) was equipped with an EM-CCD camera (Hamamatsu, model C9100-50) running MetaMorph (Universal Imaging) for data acquisition and experimental control. To accommodate the NMR system, the inverted microscope was placed beside the magnet and a 380 mm-long objective inverter (LSMtech, model 500) was used to place the objective over the chamber looking through a 650 *µ*m gap in the RF coil (Fig. 1). The upper arm of the microscope which carries the condenser, as well as the microscope stage were removed to allow access for connecting the objective inverter and the Rotation Immobilizer (to prevent the rotation of the nosepiece) (LSMtech). A 10x water immersion objective was used. Images were set to acquire every 30 s to be similar to the NMR acquisition rate.

The microscope interleaved measurements of the intrinsic optical signal (IOS) and intracellular calcium ([Ca^2+^]_i_) using Rhod-3 AM indicator. This used a multi LED set (Chroma Technology 89402 ET) with (Chroma Technology filters 89402X, 89402M). A multi-triggering LED illumination system (Excelitas Technologies X-cite XLED1) at 540-640nm was used for the fluorescence calcium imaging. A 10 mM stock of Rhod-3 AM dye (Thermo Fisher Scientific, Carlsbad, CA) in DMSO was diluted to 1:100 with aCSF and injected into the tissue as a bolus through a microelectrode under microscopic control. After injection, samples were let to rest for one hour prior to starting the experiment so that dye could diffuse through the tissue. For IOS imaging, transmitted near-infrared 680 nm light ^50^ from an LED (Thorlab M680F4) was used. The LED was attached to a fiberoptic threaded through a hole in the side of the chamber and illuminated the sample at 90^◦^ to the objective. In separate experiments without NMR, the fiberoptic was placed below the chamber to illuminate the sample at 180^◦^ to the objective. One or more areas or ROIs on the image were delineated and the average light intensity within each ROI was determined over a time series of images. Signal intensity was then plotted as a percent change from the baseline average (*S* − *S*_0_)*/S*_0_ *×* 100.

### Statistical analysis

All analysis was performed in MATLAB 2024b. For plotting averages (means) and standard deviations across samples, data was resampled to a 1-minute time-base and *t* = 0 was defined as the time that sucrose or KCl was added. Correlation coefficients between NMR and microscopy timeseries were analyzed, with *p* ≤ 0.01 deemed significant. One-sample t-tests were performed to determine if values were significantly different from baseline.

### 3-point method simulations and tests

Static gradient spin echo DEXSY signals were simulated with various parameters based on those used and found experimentally, and analyzed using the 3-point method. Simulations used a model for barrier-limited two-site exchange between a free compartment and a restricted (motionally averaged) compartment ^102^ with *R*_1_ relaxation during *t*_*m*_. See Supplementary Note 1 for details.

Previously-published data collected using the standard DEXR method method during OGD ^34^ was downloaded from https://github.com/nathanwilliamson/WaterExchangeMeasuresActiveTransport, sub-sampled, and analyzed using the 3-point method. See Fig. S3 caption for details.

## ACKNOWLEDGEMENTS

This research was supported [in part] by the Intramural Research Program of the National Institutes of Health (NIH). This work was partially funded by Military Traumatic Brain Injury Initiative (MTBI^2^) through the Uniformed Services University of the Health Sciences (USU), Bethesda, MD (award No. HU0001-24-2-0051). We thank MJ O’Donovan and M Falgairolle for helpful discussions regarding microscopy and the neonatal mouse spinal cord model, D Ide for sample chamber modifications, and D Szent-Györgyi for microscopy software technical support. Chat GPT-5.3 was used to edit for grammar and clarity.

## CONTRIBUTIONS

Conceptualization, RR, NHW, TXC, and PJB; Methodology, RR, NHW, TXC and NG; Investigation, RR, NHW, NG; Writing – Original Draft, NHW; Writing – Review & Editing, RR, NHW, TXC, NG, and PJB; Funding Acquisition, PJB; Supervision, PJB

## Supplementary Note 1: Simulation details

Static gradient spin echo DEXSY signals were simulated in MATLAB R2024a using a model for barrier-limited two-site exchange between a free compartment and a restricted compartment, ignoring spin-spin relaxation during encoding times but accounting for spin lattice relaxation (assuming the same rate *R*_1_ for both compartments) during the mixing time *t*_*m*_, using the following equation:

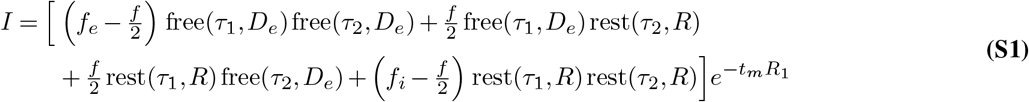

where *τ*_1_ and *τ*_2_ are the variable diffusion encoding times between the 90^◦^ and 180^◦^ pulses (i.e. half the echo time) in the first and second diffusion encoding blocks of the DEXSY sequence, *D*_*e*_ is the apparent diffusion coefficient in the free compartment, and *f*_*e*_ and *f*_*i*_ are the fractions of spins in the free and restricted compartment, respectively, with *f*_*e*_ + *f*_*i*_ = 1. Eq. S1 includes a function for attenuation due to free diffusion in the free compartment

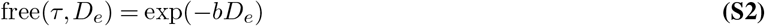

with 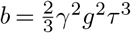, where *γ* is the gyromagnetic ratio and *g* is the gradient amplitude (set to *g* = 15.2689 as in the experimental study), and a function for attenuation due to restricted diffusion, i.e., motional averaging in a spherical compartment (Neuman, 1974),

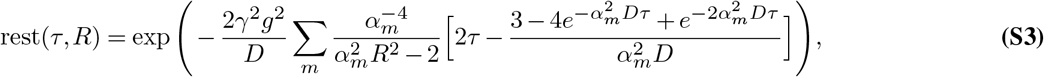

where the first five root of *α*_*m*_ = {2.0815, 5.940, 9.206, 12.405, 15.579}*/R, R* is the spherical compartment radius, and *D* is the free diffusion coefficient (2.15 × 10^−9^ m^2^/s for water at 25^◦^C).

The exchange fraction *f* was defined as

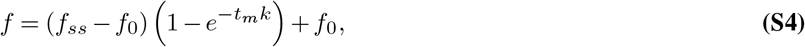

with

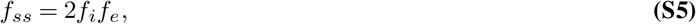

where *k* is the exchange rate constant.

Then zero-mean Gaussian noise with signal-to-noise ratio SNR was added to the signal by sampling random numbers from a normal distribution with standard deviation 1*/*SNR using randn(1, 1)*/*SNR.

Data was generated 10000 times at each ground truth *k*, with random zero-mean Gaussian noise defined by the SNR added to each signal. Ground-truth *k*’s were varied linearly over 100 values from 10−200 *s*^−1^. AXR was estimated using the 3-point method. In the base-case, the AXR was not adjusted for *R*_1_. On other cases, the AXR was adjusted for *R*_1_ by first dividing the signals by exp(−*R*_1_*t*_*m*_) using a prescribed *R*_1_, which in some cases was not the true *R*_1_. Experimentally, this is done post-hoc by utilizing prior knowledge of the apparent *R*_1_ measured separately.

## Supplementary figures

**Fig. S1.**
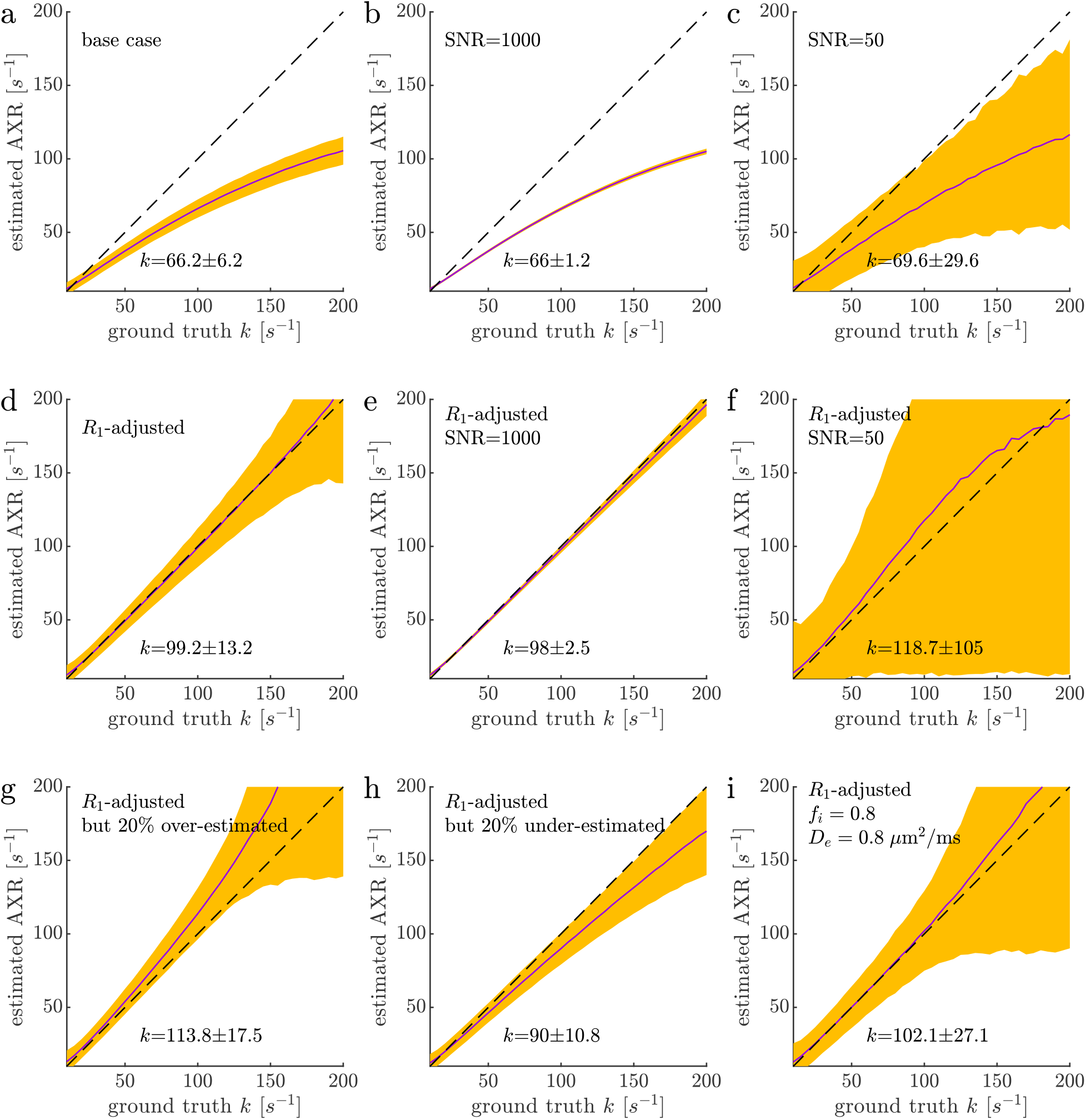
Simulation tests of accuracy and precision of AXR estimates using the 3-point method, with *t*_*m*,int_ = 10 ms. DEXSY data was simulated using Eq. S1 with radius *R* = 800 nm and a free compartment with *D*_*e*_ = 1 *µ*m^2^*/*ms. Simulations used parameters *t*_*m*_ = [0.2, 10, 160] ms and (*τ*_1_, *τ*_2_) = (0.593, 0.58) ms i.e., (*b*_1_, *b*_2_) = (2.320, 2.170) ms*/µ*m^2^, mimicking values used experimentally (see Methods). In the “base case”, *R*_1_ = 1.5 *s*^−1^, SNR = 200, and *f*_*i*_ = 0.5. Additionally, *R*_1_ was not accounted for. In other simulations, specific parameters were varied as specified in the graphs while keeping other parameters the same as in the base case. This includes adjusting for *R*_1_, either using the true value, or a value which was 20% over or under-estimated. The diagonal dashed line shows perfect accuracy. The solid purple line and orange band show the mean and standard deviation of AXR estimates as a function of the ground truth *k*. Values are displayed in each plot for the mean *±* standard deviation of AXR estimates at a ground-truth *k* = 100 *s*^−1^. Standard deviations vary with SNR as expected. In a–c, Values become more and more under-estimated as the ground-truth *k* is increased due to the the effect of spin-lattice relaxation. in d–f, the AXR estimates were adjust for *R*_1_ relaxation. This removes the under-estimation bias seen in a–c. g) The *R*_1_ value used to adjust for relaxation is 20% greater than the true *R*_1_, leading to *k* being over-estimated. h) The *R*_1_ value used to adjust for relaxation is 20% less than the true *R*_1_, leading to *k* being under-estimated. I) Relaxation is accounted for using the true *R*_1_ and parameters are the same as the base case except *f*_*i*_ = 0.8 and *D*_*e*_ = 0.8 *µ*m^2^*/*ms. Compared to (d), this leads to greater standard deviations but does not affect the accuracy of the measurement.

**Fig. S2.**
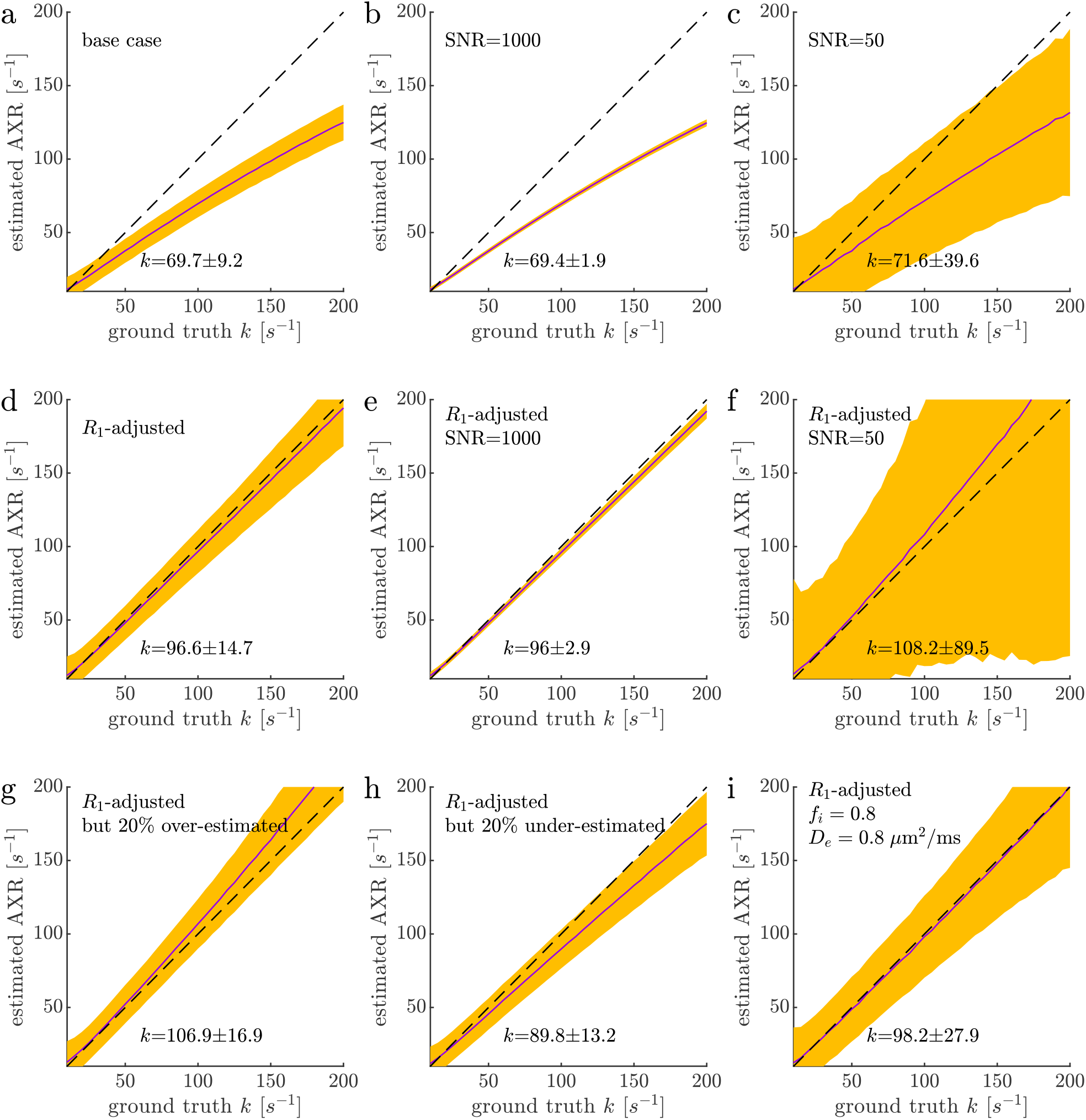
Simulation tests of accuracy and precision of AXR estimates using the 3-point method, with *t*_*m*,int_ = 5 ms. All parameters are the same as in Fig. S1, except that the intermediate mixing time was set to 5 ms rather than 10 ms. This leads to reduced precision and accuracy when the ground truth *k <* 100 s^−1^, but greater precision and accuracy for ground truth *k >* 100 s^−1^.

**Fig. S3.**
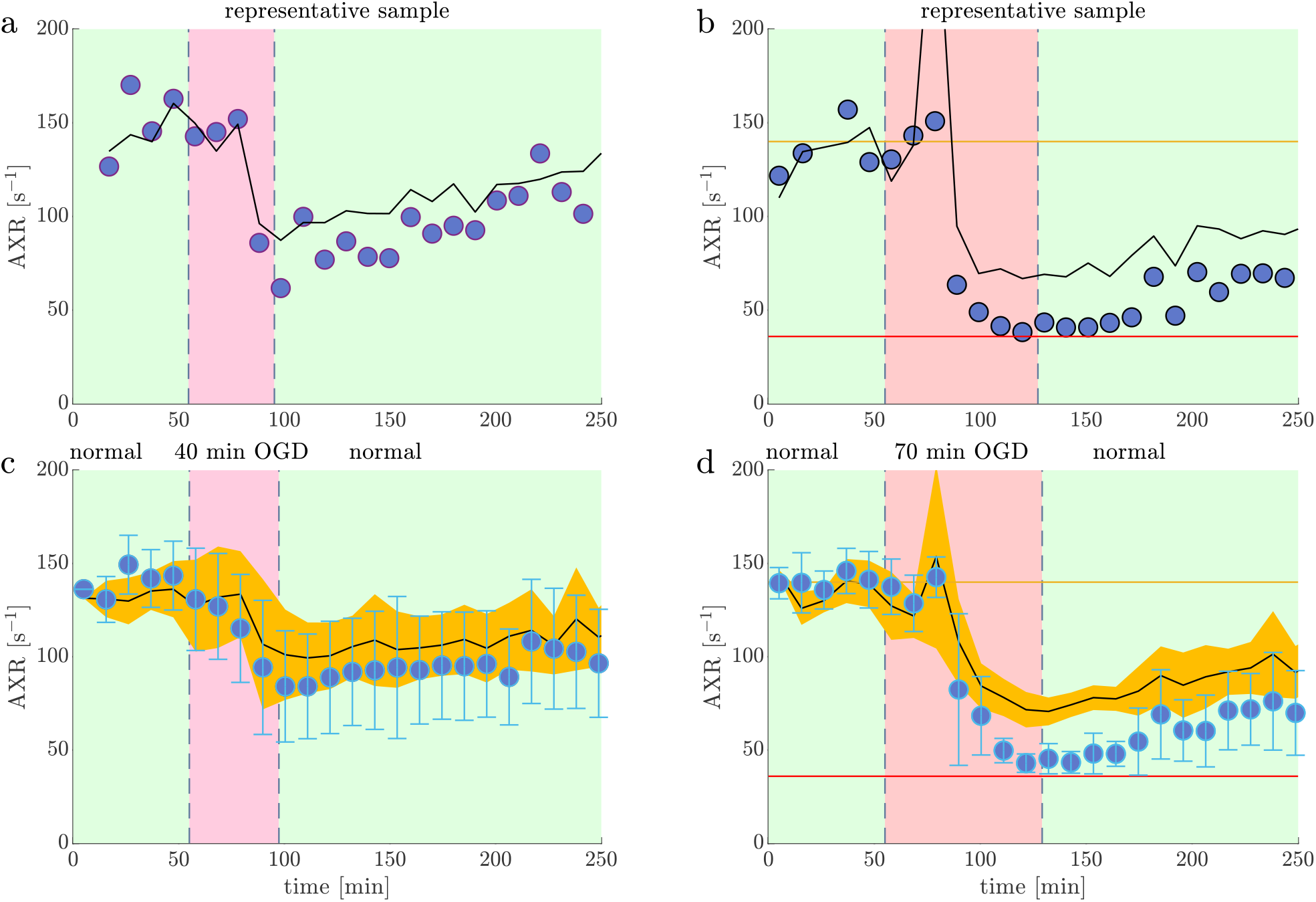
Tests of 3-point analysis by subsampling OGD data. a–d) Comparisons between the 3-point AXR analysis (solid black line) and the full DEXR method (blue circles) for representative samples (a,b) and means and standard deviations (orange bands and error bars) across all samples (c,d) from previously-published (Williamson & Ravin et al., 2023) data collected during experiments involving 40 min of oxygen and glucose deprivation (OGD) (a,c, n=8), and during 70 min OGD (b,d, n=9). The full DEXR method used 22 points from (*b*_1_, *b*_2_) = (0.089, 4.417) and (2.320, 2.170) ms*/µ*m^2^ with *t*_*m*_ = [0.2, 1, 2, 4, 7, 10, 20, 40, 80, 160, 300] ms. Signal acquired with the first (*b*_1_, *b*_2_) pair was used to calculate the apparent *R*_1_ and then remove it from signal acquired with the second (*b*_1_, *b*_2_) pair so that exchange contrast could be isolated. The 3-point analysis used the (2.320, 2.170) ms*/µ*m^2^ pair with *t*_*m*_ = [0.2, 10, 160] ms and adjusted for relaxation using the effective *R*_1_ from the full DEXR method averaged over the first 3 measurement sets in the normal condition. The 3-point analysis agrees with the full DEXR method when *k* values are near 150 s^−1^, but are biased towards higher values when *k* values decrease. The bias is not due to changes in relaxation because it is still present when the apparent *R*_1_ from each set is used for adjustment. While both models assume a first order kinetic model, the 3-point method relies more heavily on this assumption because it assumes that the important timescale for exchange is captured completely by the intermediate (10 ms) encoding time. Therefore the bias may be due to multiexponential character or deviations from a first order kinetic model.

**Fig. S4.**
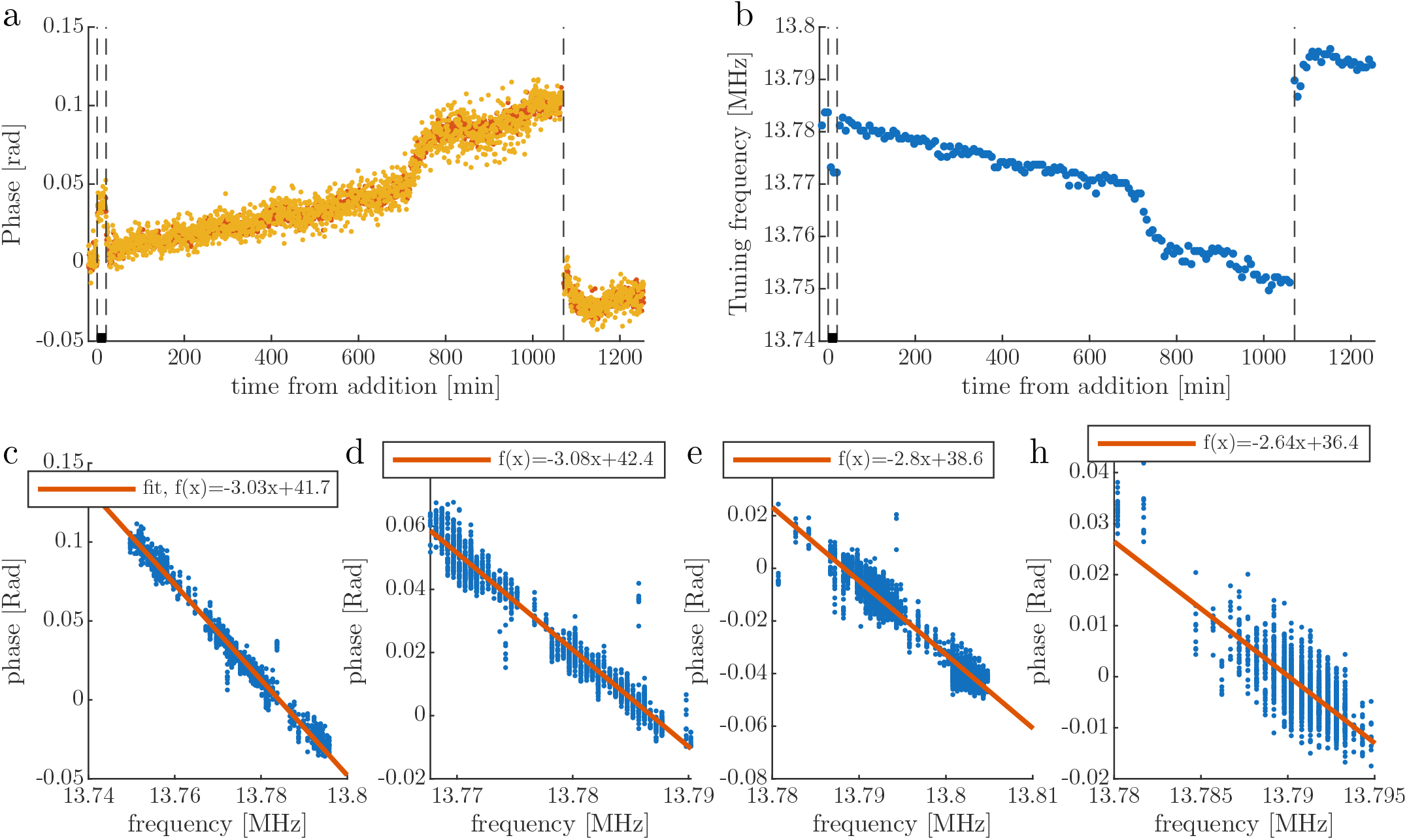
Corelation of phase vs. tuning frequency. A summary of the results of the phase and tuning measurements on pure aCSF perturbed with +50 mM KCl from t=0, then washed back to normal aCSF at 20 minutes, and then continuing to record overnight. a–b) Representative recordings during one experiment. The coil was tuned again at 1050 min. c) Correlation from that experiment. The tuning frequency showed a linear relationship with the phase. d-f) Correlations between phase and frequency for three other identical experiments. Note that the slope is similar between experiments. This indicates that the phase measures the tuning frequency.

**Fig. S5.**
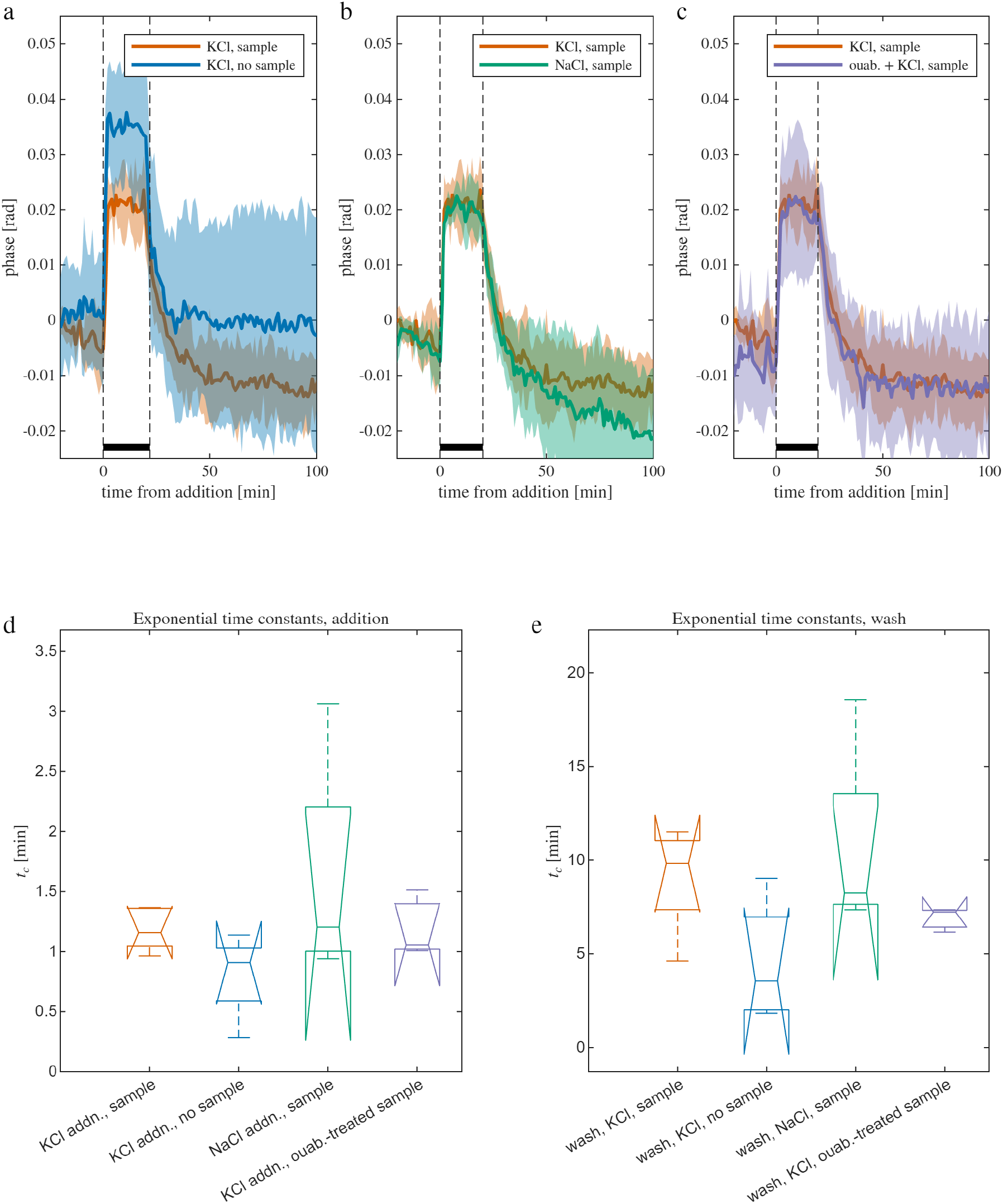
Timecourse and time constants of phase during addition and wash. a-c) Timecourses comparing between addition/wash of 50 mM KCl with a sample (n=5) to (a) 50 mM KCl without a sample (n=4), (b) 50 mM NaCl with a sample (n=4), and (c) 50 mM KCl added to a sample 1 hr after treating it with 100 *µ*M of the Na^+^*/*K^+^–ATPase, ouabain (n=3). Phase shows characteristics of equilibration, rising to a plateau and decaying towards baseline. The increase in phase is 0.018 *±* 0.004, 0.031 *±* 0.010, 0.018 *±* 0.003, and 0.018 *±* 0.010 radians when 50 mM KCl is added to a sample, 50 mM KCl is added without a sample, 50 mM NaCl is added to a sample, and 50 mM KCl is added to a sample treated with ouabain, respectively, not significantly different between groups (one-way ANOVA). Exponential time constants are 1.2 *±* 0.2 min when adding 50 mM KCl to a sample and 9.0 *±* 2.7 min when washing washing to normal aCSF. Time constants are not significantly different when no sample is present, when NaCl is added instead of KCl, and when KCl is added approximately 1 hr after 100 *µ*M of the Na^+^*/*K^+^–ATPase, ouabain was added earlier in the experiment (one-way ANOVA).

**Fig. S6.**
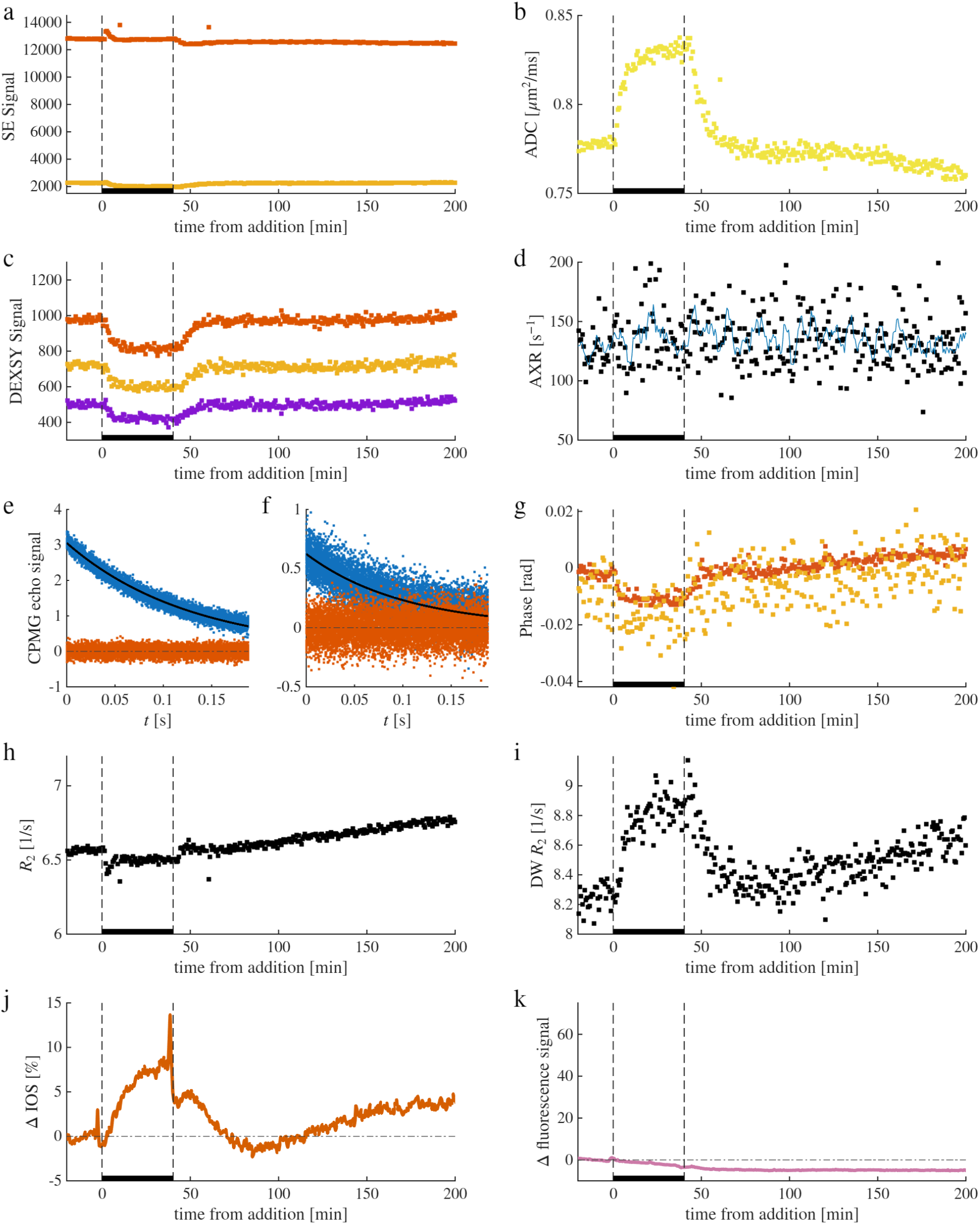
Example realtime NMR and microscopy recording during an experiment involving 100 mM sucrose (osmolyte) addition and washout. Realtime recordings of two (*b* = 0.025 and 2.25 ms*/µ*m^2^, red and orange dots, respectively) raw spin echo diffusion signals (a), processed into one ADC measurement using Eq. 1 (b), and three (*t*_*m*_ = 0.02, 10, and 160 ms, red, orange, and blue dots, respectively) raw exchange-weighted DEXSY signals (c), processed into one AXR measurement using Eq. 2 (d). In (d), a 6-point moving average of AXR is also shown (solid blue line). (e,f) Example real (blue) and imaginary (red) components of the CPMG echo signals and exponential fits (solid black line) used to estimate *R*_2_ acquired 18 minutes prior to KCl addition using the spin echo sequence with (e) *b* = 0.025 and (f) 2.25 ms*/µ*m^2^. Dot-dash line shows zero signal (g) Mean echo phase recorded the spin echo sequence with *b* = 0.025 and 2.25 ms*/µ*m^2^ (red and orange dots, respectively). (h,i) *R*_2_ estimated from the CMPG decay with the spin echo echo sequence at (h) *b* = 0.025 and (i) 2.25 ms*/µ*m^2^. Intrinsic optical signal images (j) and fluorescence images from the channel used for recording intracellular calcium from the fluorescent indicator, Rhod-3 (k) are acquired simultaneously with the NMR, and the percent change in image intensity from a region of interest (ROI) are displayed. Here, Rhod-3 was not injected, so (k) shows the change in autofluorescence at that wavelength. The lack of change indicates that autofluorescence does not confound intracellular calcium imaging. Spuriously higher signals (a) and lower *R*_2_ (h) at roughly 10 and 60 min show when tuning was performed.

**Fig. S7.**
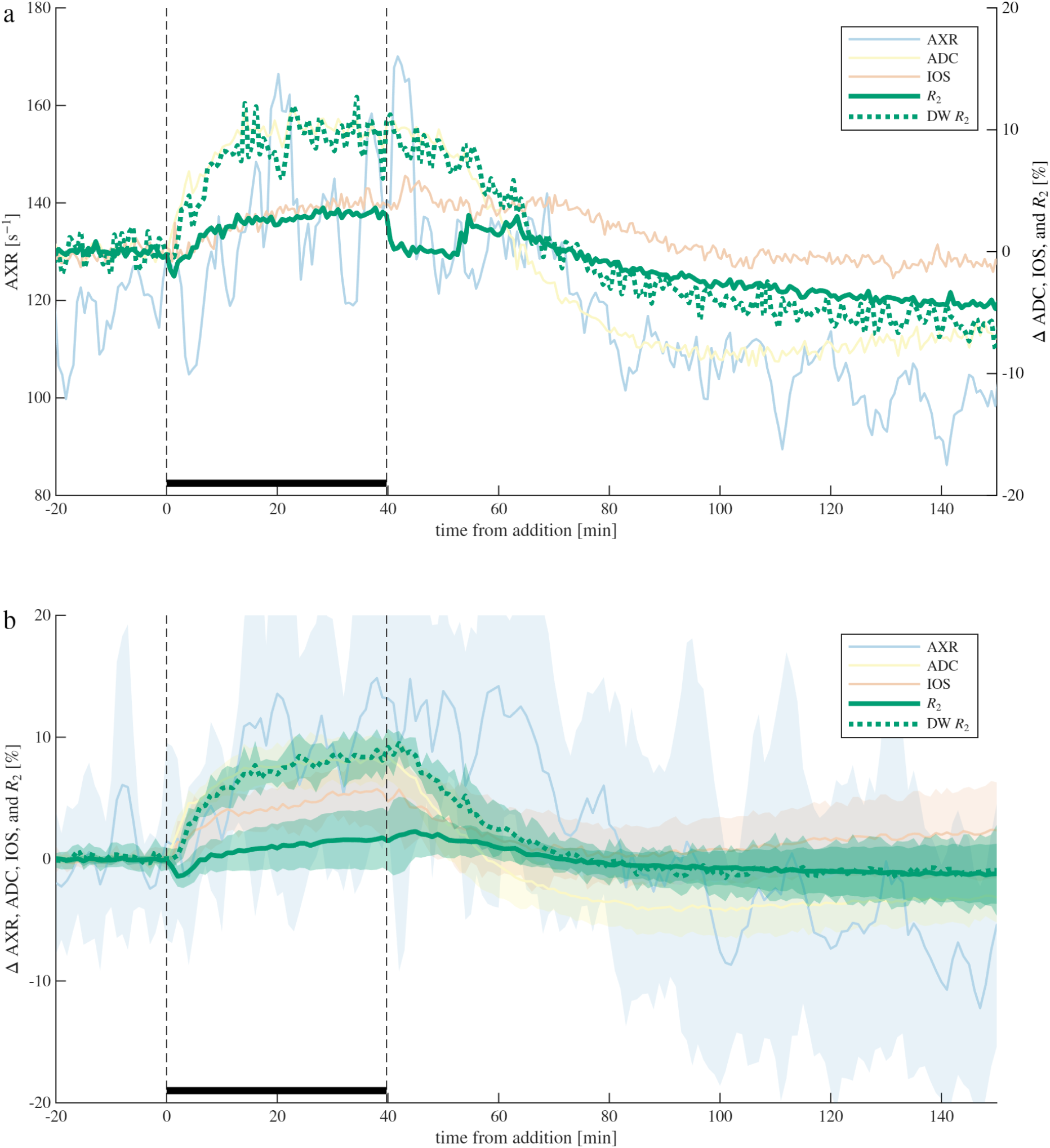
*R*_2_ and DW *R*_2_ show different trends during sucrose addition. a) Representative sample and b) mean ± SD of *R*_2_ and DW *R*_2_ plotted on top of AXR, ADC and IOS during 40 min 100 mM sucrose exposure and washout (*n* = 8).

**Fig. S8.**
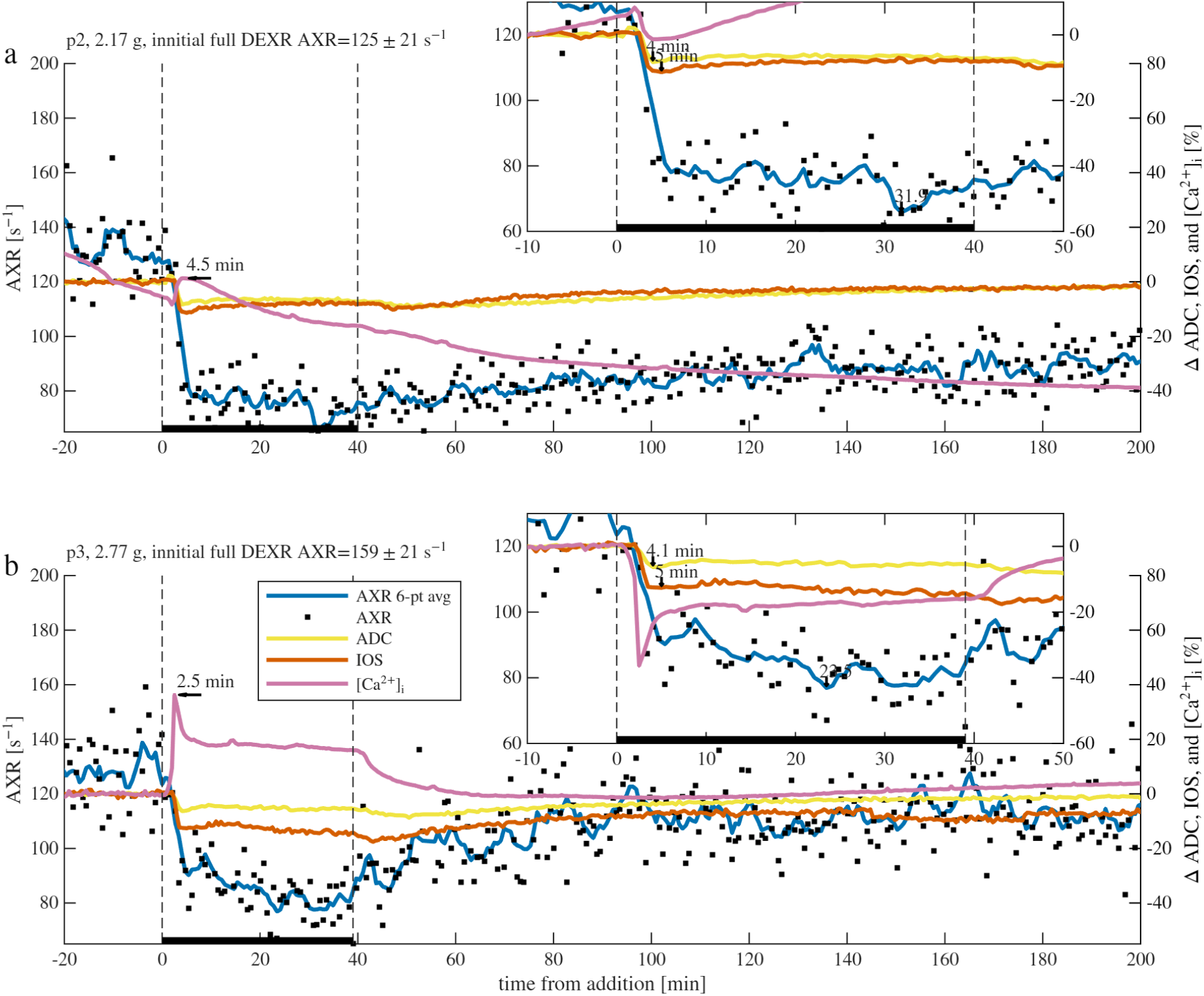
Individual recordings showing cellular shrinkage caused by 40 minutes of 50 mM KCl. a-d) Individual recordings from *n* = 2 samples. A 6-point moving average of AXR is also shown (solid blue line). Correlations were deemed significant when *p*-values were less than 0.01 and not significant (NS) otherwise. Correlation coefficients (and *p*-values) between AXR and ADC, AXR and IOS, AXR and [Ca^2+^]_i_, ADC and IOS, ADC and [Ca^2+^]_i_, and IOS and [Ca^2+^]_i_ from data acquired between t=0 to 80 minutes for each recording are [a) 0.77, 0.74, NS, 0.71, 0.50, NS (*p* ≪ 0.001, ≪ 0.001, = 0.02, ≪ 0.001, ≪ 0.001, 0.08)], [b) 0.32, 0.69, −0.67, 0.85, NS, NS (*p* = 0.003, ≪ 0.001, ≪ 0.001, ≪ 0.001, = 0.08, = 0.05)].

**Fig. S9.**
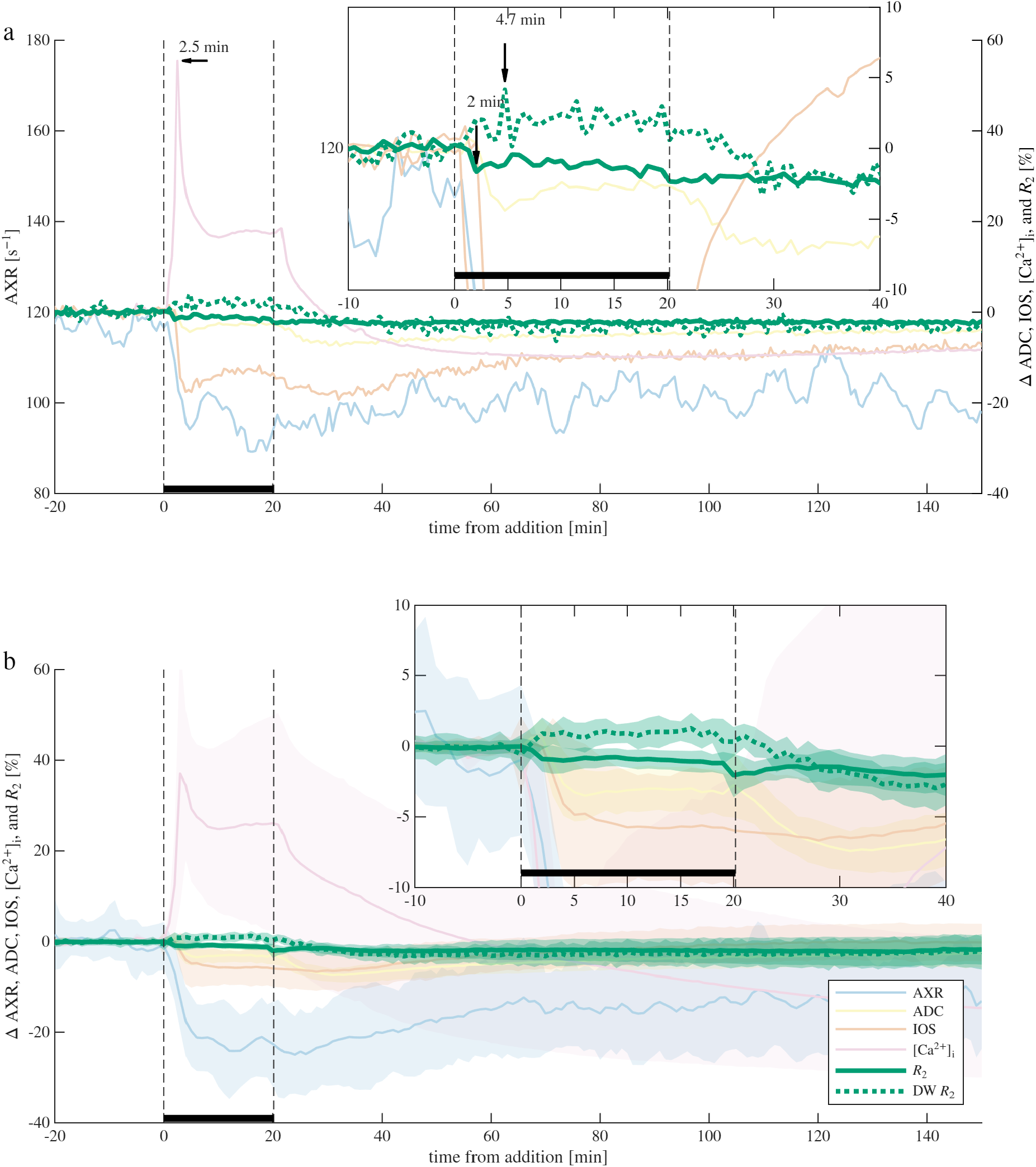
*R*_2_ and DW *R*_2_ show opposite trends during KCl addition. a) Representative sample and b) mean ± SD of *R*_2_ and DW *R*_2_ plotted on top of AXR, ADC, IOS, and [Ca^2+^]_i_ during 20 min KCl exposure and washout (*n* = 16, three of which included [Ca^2+^]_i_ imaging). Insets show zoomed regions with [Ca^2+^]_i_ inverted for comparison. Arrows in (a) mark signal minima and maxima.

## Bibliography

1. Ileana O Jelescu, Marco Palombo, Francesca Bagnato, and Kurt G Schilling. Challenges for biophysical modeling of microstructure. Journal of Neuroscience Methods, 344:108861, 2020.

2. Ileana O Jelescu, Magdalena Zurek, Kerryanne V Winters, Jelle Veraart, Anjali Rajaratnam, Nathanael S Kim, James S Babb, Timothy M Shepherd, Dmitry S Novikov, Sungheon G Kim, et al. In vivo quantification of demyelination and recovery using compartment-specific diffusion mri metrics validated by electron microscopy. Neuroimage, 132:104–114, 2016.

3. Dan Benjamini, Diego Iacono, Michal E Komlosh, Daniel P Perl, David L Brody, and Peter J Basser. Diffuse axonal injury has a characteristic multidimensional mri signature in the human brain. Brain, 144(3):800–816, 2021.

4. Alexandru V Avram, Kadharbatcha S Saleem, Michal E Komlosh, Cecil C Yen, Frank Q Ye, and Peter J Basser. High-resolution cortical map-mri reveals areal borders and laminar substructures observed with histological staining. NeuroImage, 264:119653, 2022.

5. Amy FD Howard, Istvan N Huszar, Adele Smart, Michiel Cottaar, Greg Daubney, Taylor Hanayik, Alexandre A Khrapitchev, Rogier B Mars, Jeroen Mollink, Connor Scott, et al. An open resource combining multi-contrast mri and microscopy in the macaque brain. Nature communications, 14(1):4320, 2023.

6. Kurt G Schilling, Francesco Grussu, Andrada Ianus, Brian Hansen, Amy FD Howard, Rachel LC Barrett, Manisha Aggarwal, Stijn Michielse, Fatima Nasrallah, Warda Syeda, et al. Considerations and recommendations from the ismrm diffusion study group for pre-clinical diffusion mri: Part 2—ex vivo imaging: Added value and acquisition. Magnetic resonance in medicine, 93(6):2535–2560, 2025.

7. Timothy M Shepherd, Peter E Thelwall, Greg J Stanisz, and Stephen J Blackband. Aldehyde fixative solutions alter the water relaxation and diffusion properties of nervous tissue. Magnetic Resonance in Medicine: An Official Journal of the International Society for Magnetic Resonance in Medicine, 62(1):26–34, 2009.

8. Nathan H Williamson, Rea Ravin, Dan Benjamini, Hellmut Merkle, Melanie Falgairolle, Michael James O’Donovan, Dvir Blivis, Dave Ide, Teddy X Cai, Nima S Ghorashi, et al. Magnetic resonance measurements of cellular and sub-cellular membrane structures in live and fixed neural tissue. Elife, 8:e51101, 2019.

9. Helene Benveniste, Laurence W Hedlund, and G Allan Johnson. Mechanism of detection of acute cerebral ischemia in rats by diffusion-weighted magnetic resonance microscopy. Stroke, 23(5):746–754, 1992.

10. Lawrence L Latour, Yasuhiro Hasegawa, James E Formato, Marc Fisher, and Christopher H Sotak. Spreading waves of decreased diffusion coefficient after cortical stimulation in the rat brain. Magnetic resonance in medicine, 32(2):189–198, 1994.

11. Denis Le Bihan, Shin-ichi Urayama, Toshihiko Aso, Takashi Hanakawa, and Hidenao Fukuyama. Direct and fast detection of neuronal activation in the human brain with diffusion mri. Proceedings of the National Academy of Sciences, 103(21):8263–8268, 2006.

12. Daniel Nunes, Andrada Ianus, and Noam Shemesh. Layer-specific connectivity revealed by diffusion-weighted functional mri in the rat thalamocortical pathway. Neuroimage, 184: 646–657, 2019.

13. Inès de Riedmatten, Arthur PC Spencer, Wiktor Olszowy, and Ileana O Jelescu. Apparent diffusion coefficient fmri shines light on white matter resting-state connectivity compared to bold. Communications biology, 8(1):447, 2025.

14. Andreea Hertanu, Tommaso Pavan, and Ileana O. Jelescu. Somatosensory-evoked response induces extensive diffusivity and kurtosis changes associated with neural activity in rodents. Imaging Neuroscience, 3:00445, 02 2025. ISSN 2837-6056. doi: 10.1162/imag_a_00445.

15. Karla L Miller, Daniel P Bulte, Hannah Devlin, Matthew D Robson, Richard G Wise, Mark W Woolrich, Peter Jezzard, and Timothy EJ Behrens. Evidence for a vascular contribution to diffusion fmri at high b value. Proceedings of the National Academy of Sciences, 104(52): 20967–20972, 2007.

16. Kulam Najmudeen Magdoom, Alexandru V Avram, Joelle E Sarlls, and Peter J Basser. In vivo palpation of anisotropic human brain tissue using mri. bioRxiv, pages 2025–06, 2025.

17. Elmar Busch, Mathias Hoehn-Berlage, Manfred Eis, Michael L Gyngell, and Konstantin-Alexander Hossmann. Simultaneous recording of eeg, dc potential and diffusion-weighted nmr imaging during potassium induced cortical spreading depression in rats. NMR in Biomedicine, 8(2):59–64, 1995.

18. Meng Cui, Yifeng Zhou, Bowen Wei, Xiao-Hong Zhu, Wei Zhu, Mark A Sanders, Kamil Ugurbil, and Wei Chen. A proof-of-concept study for developing integrated two-photon microscopic and magnetic resonance imaging modality at ultrahigh field of 16.4 tesla. Scientific reports, 7(1):2733, 2017.

19. Evelyn MR Lake, Xinxin Ge, Xilin Shen, Peter Herman, Fahmeed Hyder, Jessica A Cardin, Michael J Higley, Dustin Scheinost, Xenophon Papademetris, Michael C Crair, et al. Simultaneous cortex-wide fluorescence ca2+ imaging and whole-brain fmri. Nature methods, 17 (12):1262–1271, 2020.

20. Jörg Kärger, Harry Pfeifer, and Wilfried Heink. Principles and application of self-diffusion measurements by nuclear magnetic resonance. In Advances in Magnetic and optical resonance, volume 12, pages 1–89. Elsevier, 1988.

21. P.T. Callaghan. Translational Dynamics and Magnetic Resonance Principles of Pulsed Gradient Spin Echo NMR. Oxford University Press, 2011.

22. P. T. Callaghan and I. Furó. Diffusion-diffusion correlation and exchange as a signature for local order and dynamics. The Journal of Chemical Physics, 120(8):4032–4038, 2004.

23. Ingrid Åslund, Agnieszka Nowacka, Markus Nilsson, and Daniel Topgaard. Filter-exchange PGSE NMR determination of cell membrane permeability. Journal of Magnetic Resonance, 200(2):291–295, 2009. ISSN 1090-7807.

24. Xin Tian, Hua Li, Xiaoyu Jiang, Jingping Xie, John C Gore, and Junzhong Xu. Evaluation and comparison of diffusion mr methods for measuring apparent transcytolemmal water exchange rate constant. Journal of Magnetic Resonance, 275:29–37, 2017.

25. Dominik Ludwig, Frederik Bernd Laun, Mark Edward Ladd, Peter Bachert, and Tristan Anselm Kuder. Apparent exchange rate imaging: On its applicability and the connection to the real exchange rate. Magnetic Resonance in Medicine, 86(2):677–692, 2021.

26. Mohammad Khateri, Marco Reisert, Alejandra Sierra, Jussi Tohka, and Valerij G Kiselev. What does fexi measure? NMR in Biomedicine, 35(12):e4804, 2022.

27. Valerij G Kiselev and Jing-Rebecca Li. What does fexi measure in neurons? arXiv preprint arXiv:2601.20657, 2026.

28. Andrada Ianus, Daniel C Alexander, Hui Zhang, and Marco Palombo. Mapping complex cell morphology in the grey matter with double diffusion encoding mr: A simulation study. Neuroimage, 241:118424, 2021.

29. Arthur Chakwizira, Kadir Ş imşek, Filip Szczepankiewicz, Marco Palombo, and Markus Nilsson. The role of dendritic spines in water exchange measurements with diffusion mri: Double diffusion encoding and free-waveform mri. NMR in Biomedicine, 39(7):e70315, 2026.

30. Kadir Ş imşek, Arthur Chakwizira, Markus Nilsson, and Marco Palombo. The role of dendritic spines in water exchange measurements with diffusion mri: Time-dependent single diffusion encoding mri. arXiv preprint arXiv:2506.18229, 2025.

31. Teddy X Cai, Nathan H Williamson, and Peter J Basser. Localization-driven exchange contrast in diffusion exchange spectroscopy. Journal of Magnetic Resonance, 390:108118, 2026.

32. Nathan H Williamson, Rea Ravin, Teddy X Cai, Julian A Rey, and Peter J Basser. Hydrophysiology nmr reveals mechanisms of steady-state water exchange in neural tissue. bioRxiv, pages 2024–12, 2024.

33. Nathan H Williamson, Rea Ravin, Teddy X Cai, Julian A Rey, and Peter J Basser. Passive water exchange between multiple sites can explain why apparent exchange rate constants depend on ionic and osmotic conditions in gray matter. Magnetic Resonance Letters, page 200225, 2025. ISSN 2772-5162. doi: 10.1016/j.mrl.2025.200225.

34. Nathan H Williamson, Rea Ravin, Teddy X Cai, Melanie Falgairolle, Michael J O’Donovan, and Peter J Basser. Water exchange rates measure active transport and homeostasis in neural tissue. PNAS nexus, 2(3):pgad056, 2023.

35. AS Verkman. Water permeability measurement in living cells and complex tissues. The Journal of membrane biology, 173(2):73–87, 2000.

36. Yajie Zhang, Marie Poirier-Quinot, Charles S Springer Jr, and James A Balschi. Active trans-plasma membrane water cycling in yeast is revealed by nmr. Biophysical journal, 101(11):2833–2842, 2011.

37. Ruiliang Bai, Charles S Springer Jr, Dietmar Plenz, and Peter J Basser. Fast, na+/k+ pump driven, steady-state transcytolemmal water exchange in neuronal tissue: A study of rat brain cortical cultures. Magnetic Resonance in Medicine, 79(6):3207–3217, 2018.

38. Charles S Springer Jr. Using 1h2o mr to measure and map sodium pump activity in vivo. Journal of Magnetic Resonance, 291:110–126, 2018.

39. Eva Syková and Charles Nicholson. Diffusion in brain extracellular space. Physiological reviews, 88(4):1277–1340, 2008.

40. Charlie Aird-Rossiter, Hui Zhang, Daniel C Alexander, Derek K Jones, and Marco Palombo. Decoding gray matter: large-scale analysis of brain cell morphometry to inform microstructural modeling of diffusion mr signals. arXiv preprint arXiv:2501.02100, 2025.

41. Daniel Huster, Albert J Jin, Klaus Arnold, and Klaus Gawrisch. Water permeability of polyunsaturated lipid membranes measured by 17o nmr. Biophysical journal, 73(2):855–864, 1997.

42. John C Mathai, Stephanie Tristram-Nagle, John F Nagle, and Mark L Zeidel. Structural determinants of water permeability through the lipid membrane. The Journal of general physiology, 131(1):69–76, 2008.

43. Brian A MacVicar and Daryl Hochman. Imaging of synaptically evoked intrinsic optical signals in hippocampal slices. Journal of Neuroscience, 11(5):1458–1469, 1991.

44. Knut Holthoff and Otto W Witte. Intrinsic optical signals in rat neocortical slices measured with near-infrared dark-field microscopy reveal changes in extracellular space. Journal of Neuroscience, 16(8):2740–2749, 1996.

45. PG Aitken, D Fayuk, GG Somjen, and DA Turner. Use of intrinsic optical signals to monitor physiological changes in brain tissue slices. Methods, 18(2):91–103, 1999.

46. Katharina Buchheim, Sebastian Schuchmann, Herbert Siegmund, Hans Jürgen Gabriel, Uwe Heinemann, and Hartmut Meierkord. Intrinsic optical signal measurements reveal characteristic features during different forms of spontaneous neuronal hyperactivity associated with ecs shrinkage in vitro. European Journal of Neuroscience, 11(6):1877–1882, 1999.

47. Michael Müller and George G Somjen. Intrinsic optical signals in rat hippocampal slices during hypoxia-induced spreading depression-like depolarization. Journal of neurophysiology, 82(4):1818–1831, 1999.

48. Brian A Macvicar, Denise Feighan, Angus Brown, and Bruce Ransom. Intrinsic optical signals in the rat optic nerve: role for k+ uptake via nkcc1 and swelling of astrocytes. Glia, 37(2):114–123, 2002.

49. Dmitriy Fayuk, Peter G Aitken, George G Somjen, and Dennis A Turner. Two different mechanisms underlie reversible, intrinsic optical signals in rat hippocampal slices. Journal of neurophysiology, 87(4):1924–1937, 2002.

50. Eva Syková, Lýdia Vargová, Sárka Kubinová, Pavla Jendelová, and Alexandr Chvátal. The relationship between changes in intrinsic optical signals and cell swelling in rat spinal cord slices. Neuroimage, 18(2):214–230, 2003.

51. Sonya Bahar, Minah Suh, Mingrui Zhao, and Theodore H Schwartz. Intrinsic optical signal imaging of neocortical seizures: the ‘epileptic dip’. Neuroreport, 17(5):499–503, 2006.

52. Ruiliang Bai, Andreas Klaus, Tim Bellay, Craig Stewart, Sinisa Pajevic, Uri Nevo, Hellmut Merkle, Dietmar Plenz, and Peter J Basser. Simultaneous calcium fluorescence imaging and mr of ex vivo organotypic cortical cultures: a new test bed for functional mri. NMR in Biomedicine, 28(12):1726–1738, 2015.

53. Arnold M Friedman and Joseph W Kennedy. The self-diffusion coefficients of potassium, cesium, iodide and chloride ions in aqueous solutions1. Journal of the American Chemical Society, 77(17):4499–4501, 1955.

54. Nancy Ekdawi-Sever, Juan J de Pablo, Emily Feick, and Ernst von Meerwall. Diffusion of sucrose and α, α-trehalose in aqueous solutions. The Journal of Physical Chemistry A, 107(6):936–943, 2003.

55. James A Goodman, Christopher D Kroenke, G Larry Bretthorst, Joseph JH Ackerman, and Jeffrey J Neil. Sodium ion apparent diffusion coefficient in living rat brain. Magnetic Resonance in Medicine: An Official Journal of the International Society for Magnetic Resonance in Medicine, 53(5):1040–1045, 2005.

56. George G Somjen. Mechanisms of spreading depression and hypoxic spreading depression-like depolarization. Physiological reviews, 81(3):1065–1096, 2001.

57. Jens P Dreier. The role of spreading depression, spreading depolarization and spreading ischemia in neurological disease. Nature medicine, 17(4):439–447, 2011.

58. Hiss Martins-Ferreira, Maiken Nedergaard, and Charles Nicholson. Perspectives on spreading depression. Brain research reviews, 32(1):215–234, 2000.

59. Ning Zhou, Grant RJ Gordon, Denise Feighan, and Brian A MacVicar. Transient swelling, acidification, and mitochondrial depolarization occurs in neurons but not astrocytes during spreading depression. Cerebral cortex, 20(11):2614–2624, 2010.

60. Peter J Basser. Inferring microstructural features and the physiological state of tissues from diffusion-weighted images. NMR in Biomedicine, 8(7):333–344, 1995.

61. Ying Qiao, Petrik Galvosas, Thorsteinn Adalsteinsson, Monika Schönhoff, and Paul T Callaghan. Diffusion exchange nmr spectroscopic study of dextran exchange through poly-electrolyte multilayer capsules. The Journal of chemical physics, 122(21):214912, 2005.

62. Oliver Neudert, Siegfried Stapf, and Carlos Mattea. Diffusion exchange nmr spectroscopy in inhomogeneous magnetic fields. Journal of Magnetic Resonance, 208(2):256–261, 2011.

63. Samo Lasič, Markus Nilsson, Jimmy Lätt, Freddy Ståhlberg, and Daniel Topgaard. Apparent exchange rate mapping with diffusion MRI. Magnetic Resonance in Medicine, 66(2): 356–365, 2011.

64. Dan Benjamini, Michal E. Komlosh, and Peter J. Basser. Imaging local diffusive dynamics using diffusion exchange spectroscopy MRI. Physical Review Letters, 118:158003, Apr 2017.

65. Otto Mankinen, Vladimir V Zhivonitko, Anne Selent, Sarah Mailhiot, Sanna Komulainen, Nønne L Prisle, Susanna Ahola, and Ville-Veikko Telkki. Ultrafast diffusion exchange nuclear magnetic resonance. Nature communications, 11(1):1–8, 2020.

66. Lipeng Ning, Markus Nilsson, Samo Lasič, Carl-Fredrik Westin, and Yogesh Rathi. Cumulant expansions for measuring water exchange using diffusion mri. The Journal of chemical physics, 148(7):074109, 2018.

67. Teddy X Cai, Dan Benjamini, Michal E Komlosh, Peter J Basser, and Nathan H Williamson. Rapid detection of the presence of diffusion exchange. Journal of Magnetic Resonance, 297:17–22, 2018.

68. Nathan H Williamson, Rea Ravin, Teddy X Cai, Dan Benjamini, Melanie Falgairolle, Michael J O’Donovan, and Peter J Basser. Real-time measurement of diffusion exchange rate in biological tissue. Journal of Magnetic Resonance, 317:106782, 2020.

69. Yuval Scher, Shlomi Reuveni, and Yoram Cohen. Constant gradient fexsy: A time-efficient method for measuring exchange. Journal of Magnetic Resonance, 311:106667, 2020.

70. Teddy X Cai, Nathan H Williamson, Rea Ravin, and Peter J Basser. Disentangling the effects of restriction and exchange with diffusion exchange spectroscopy. Frontiers in Physics, page 223, 2022.

71. Arthur Chakwizira, Carl-Fredrik Westin, Jan Brabec, Samo Lasič, Linda Knutsson, Filip Szczepankiewicz, and Markus Nilsson. Diffusion mri with pulsed and free gradient waveforms: effects of restricted diffusion and exchange. NMR in Biomedicine, page e4827, 2022.

72. Teddy X Cai, Nathan H Williamson, Rea Ravin, and Peter J Basser. The diffusion exchange ratio (dexr): A minimal sampling of diffusion exchange spectroscopy to probe exchange, restriction, and time-dependence. Journal of Magnetic Resonance, page 107745, 2024.

73. D.G. Rata, F. Casanova, J. Perlo, D.E. Demco, and B. Blümich. Self-diffusion measurements by a mobile single-sided nmr sensor with improved magnetic field gradient. Journal of Magnetic Resonance, 180(2):229–235, 2006. ISSN 1090-7807. doi: 10.1016/j.jmr.2006.02.015.

74. Teddy Xuke Cai. In search of lost time: development of rapid magnetic resonance methods to probe time-varying diffusion. PhD thesis, University of Oxford, 2023.

75. Herman Y Carr and Edward M Purcell. Effects of diffusion on free precession in nuclear magnetic resonance experiments. Physical review, 94(3):630, 1954.

76. Saul Meiboom and David Gill. Modified spin-echo method for measuring nuclear relaxation times. Review of scientific instruments, 29(8):688–691, 1958.

77. Federico Casanova, Juan Perlo, and Bernhard Blümich. Single-sided NMR. Springer, 2011.

78. Teddy Xuke Cai, N Williamson, Rea Ravin, and Peter Joel Basser. Multiexponential analysis of diffusion exchange times reveals a distinct exchange process associated with metabolic activity. In Proceedings of the 32nd Annual Meeting of ISMRM, Toronto, Canada, 2023.

79. Alfredo Ordinola, Evren Özarslan, Ruiliang Bai, and Magnus Herberthson. Limitations and generalizations of the first order kinetics reaction expression for modeling diffusion-driven exchange: Implications on nmr exchange measurements. The Journal of Chemical Physics, 160(8), 2024.

80. Giulio Giovannetti, Francesca Frijia, Luca Menichetti, Valentina Hartwig, Vittorio Viti, and Luigi Landini. An efficient method for electrical conductivity measurement in the rf range. Concepts in Magnetic Resonance Part B: Magnetic Resonance Engineering, 37(3):160–166, 2010.

81. Manushka V Vaidya, Christopher M Collins, Daniel K Sodickson, Ryan Brown, Graham C Wiggins, and Riccardo Lattanzi. Dependence of and field patterns of surface coils on the electrical properties of the sample and the mr operating frequency. Concepts in Magnetic Resonance Part B: Magnetic Resonance Engineering, 46(1):25–40, 2016.

82. Paul T Callaghan and Yang Xia. Velocity and diffusion imaging in dynamic nmr microscopy. Journal of Magnetic Resonance (1969), 91(2):326–352, 1991.

83. N. Bloembergen, E. M. Purcell, and R. V. Pound. Relaxation effects in nuclear magnetic resonance absorption. Phys. Rev., 73:679–712, Apr 1948. doi: 10.1103/PhysRev.73.679.

84. Ryogo Kubo. A stochastic theory of line shape. Advances in chemical physics, 15:101–127, 1969.

85. Kenneth R Brownstein and CE Tarr. Importance of classical diffusion in nmr studies of water in biological cells. Physical review A, 19(6):2446, 1979.

86. Thomas J Jentsch. Vracs and other ion channels and transporters in the regulation of cell volume and beyond. Nature Reviews Molecular Cell Biology, 17(5):293, 2016.

87. Botros Shenoda. The role of na+/ca2+ exchanger subtypes in neuronal ischemic injury. Translational stroke research, 6(3):181–190, 2015.

88. Rune Enger, Wannan Tang, Gry Fluge Vindedal, Vidar Jensen, P Johannes Helm, Rolf Sprengel, Loren L Looger, and Erlend A Nagelhus. Dynamics of ionic shifts in cortical spreading depression. Cerebral Cortex, 25(11):4469–4476, 2015.

89. Matthew A Stern, Eric R Cole, Claire-Anne Gutekunst, Jenny J Yang, Ken Berglund, and Robert E Gross. Organellular imaging in vivo reveals a depletion of endoplasmic reticular calcium during post-ictal cortical spreading depolarization. bioRxiv, 2024.

90. Trent A Basarsky, Steven N Duffy, R David Andrew, and Brian A MacVicar. Imaging spreading depression and associated intracellular calcium waves in brain slices. Journal of Neuroscience, 18(18):7189–7199, 1998.

91. Jessica L Seidel, Carole Escartin, Cenk Ayata, Gilles Bonvento, and C William Shuttleworth. Multifaceted roles for astrocytes in spreading depolarization: A target for limiting spreading depolarization in acute brain injury? Glia, 64(1):5–20, 2016.

92. Nathan H Williamson, Velencia J Witherspoon, Teddy X Cai, Rea Ravin, Ferenc Horkay, and Peter J Basser. Low-field, high-gradient nmr shows diffusion contrast consistent with localization or motional averaging of water near surfaces. Magnetic Resonance Letters, 2023.

93. Michael E Moseley, Y Cohen, J Mintorovitch, L Chileuitt, H Shimizu, J Kucharczyk, MF Wendland, and PR Weinstein. Early detection of regional cerebral ischemia in cats: comparison of diffusion-and t2-weighted mri and spectroscopy. Magnetic resonance in medicine, 14(2):330–346, 1990.

94. Adam W Anderson, Jianhui Zhong, Ognen AC Petroff, Aaron Szafer, Bruce R Ransom, James W Prichard, and John C Gore. Effects of osmotically driven cell volume changes on diffusion-weighted imaging of the rat optic nerve. Magnetic resonance in medicine, 35 (2):162–167, 1996.

95. David L Buckley, Jonathan D Bui, M Ian Phillips, Tibor Zelles, Benjamin A Inglis, H Daniel Plant, and Stephen J Blackband. The effect of ouabain on water diffusion in the rat hippocampal slice measured by high resolution nmr imaging. Magnetic Resonance in Medicine: An Official Journal of the International Society for Magnetic Resonance in Medicine, 41(1):137–142, 1999.

96. AW Anderson, J Xie, J Pizzonia, RA Bronen, DD Spencer, and JC Gore. Effects of cell volume fraction changes on apparent diffusion in human cells. Magnetic resonance imaging, 18(6):689–695, 2000.

97. Ileana Ozana Jelescu, Luisa Ciobanu, Françoise Geffroy, Pierre Marquet, and Denis Le Bihan. Effects of hypotonic stress and ouabain on the apparent diffusion coefficient of water at cellular and tissue levels in aplysia. NMR in Biomedicine, 27(3):280–290, 2014.

98. Yuval Scher, Shlomi Reuveni, and Yoram Cohen. Quantifying transmembrane water exchange by diffusion nmr methods: From yeast cells to optic nerve ex vivo. NMR in Biomedicine, 39(3):e70224, 2026.

99. Nathan H Williamson, Rea Ravin, Teddy X Cai, Julian A Rey, and Peter J Basser. Steady-state water exchange in neural tissue is primarily passive and through the phospholipid bilayer. bioRxiv, pages 2024–12, 2024.

100. G Eidmann, R Savelsberg, Peter Blümler, and Bernhard Blümich. The nmr mouse, a mobile universal surface explorer. Journal of Magnetic Resonance, 122:104–109, 1996.

101. Erwin L Hahn. Spin echoes. Physical review, 80(4):580, 1950.

102. CH Neuman. Spin echo of spins diffusing in a bounded medium. The Journal of Chemical Physics, 60(11):4508–4511, 1974.

